# A phased chromosome-level genome assembly provides insights into the evolution of sex chromosomes in *Amaranthus tuberculatus*

**DOI:** 10.1101/2024.05.30.596720

**Authors:** Damilola A. Raiyemo, Luan Cutti, Eric L. Patterson, Victor Llaca, Kevin Fengler, Jacob S. Montgomery, Sarah Morran, Todd A. Gaines, Patrick J. Tranel

## Abstract

- *Amaranthus tuberculatus* (waterhemp) is a troublesome weed species of agronomic importance that is dioecious with an XY sex-determination system. The evolution of sex chromosomes, the contiguity of sex-determining region (SDR) and the expression pattern of genes within the SDR remain poorly understood.
- We assembled the genome of a male *A. tuberculatus*, phased the genome into two chromosome-level haplotypes, and performed restriction site-associated DNA genome-wide association (RAD-GWA) analysis, comparative genomics, adaptive evolution analysis, and, with existing data, transcriptomic profiling to characterize the species’ sex chromosomes.
- Comparative analysis enabled the identification of a ∼32.8 Mb SDR on chromosome 1 that is gene-poor, abundant in long terminal repeat (LTR) retrotransposons, and harbors two inversions. Synteny analysis revealed that chromosome 1 likely originated from the fusion of two ancestral chromosomes, and mRNA data indicated 93 genes out of the 531 protein-coding genes within the SDR of haplome 2 were differentially expressed between mature male and female flowers, with several of the genes enriched for Gene Ontology (GO) terms involved in floral development.
- Beyond adding to our understanding of sex chromosome evolution, the genomic resource provided here will be valuable for addressing further questions on adaptive trait evolution in *Amaranthus*.

## Introduction

Dioecy, the separation of male and female reproductive systems on different plants, has evolved multiple times independently across many lineages, occurring in as many as 6% of flowering plant species (Renner & Ricklefs, 1995; Ming *et al.*, 2011; Renner, 2014), and via several mechanisms (Charlesworth & Charlesworth, 1978; Bawa, 1980; Lloyd, 1980; Henry *et al.*, 2018). One model (two-gene model) postulates that dioecy could evolve via a gynodioecy pathway requiring two mutations (Charlesworth & Charlesworth, 1978; Akagi & Charlesworth, 2019; Cronk & Müller, 2020) while the second model (one-gene model) postulates that dioecy could evolve via a single regulatory factor (Henry *et al.*, 2018). Both models have been supported by the discovery of either two [e.g., in *Actinidia* spp. (Akagi *et al.*, 2019) and *Asparagus officinalis* (Harkess *et al.*, 2020)] or one sex-determining gene(s) [e.g., in *Diospyros lotus* (Akagi *et al.*, 2014) and several *Populus* species (Müller *et al.*, 2020)].

The consequence of these models in the evolution of sex chromosome is that a sex-determining region (SDR) could evolve into a small region (e.g., ∼150 kb in *Vitis* spp.) (Massonnet *et al.*, 2020) or a larger non-recombining region that may have different evolutionary histories (i.e., strata) (e.g., 17.42 Mb in *Spinacia oleracea*) (Ma *et al.*, 2022) and have accumulated repetitive sequences due to less effective selection in low recombination regions (Na *et al.*, 2014; Hobza *et al.*, 2015; Charlesworth, 2016). The study of sex chromosome evolution has been complicated due to assembly challenges posed by gene turnovers, structural variations, and repetitive sequences common within SDRs (Charlesworth, 2019), and also the possibility of numerous floral development genes acting as the sex-determining gene(s) (Ming *et al.*, 2011). Nevertheless, the availability of long-read sequencing technologies, the ability to now detect genome-wide chromatin interactions (e.g., Hi-C), optical mapping, and overall improvements in computational approaches have made the sequencing and assembly of whole genomes, and more importantly, entire sex chromosomes, possible, thereby facilitating our understanding of sex chromosome evolution in several species (Akagi *et al.*, 2023; Du *et al.*, 2023; Healey *et al.*, 2023; Kafkas *et al.*, 2023).

*Amaranthus tuberculatus* (Moq.) J.D. Sauer (waterhemp) is a troublesome dioecious weed of agronomic crops, native to the Midwest of the United States, with a range that has expanded globally (Sauer, 1957; Steckel, 2007). It is one of nine dioecious amaranths in the subgenus *Acnida* (L.) (Sauer, 1957, 1972; Mosyakin & Robertson, 1996). Due to the failure of several important herbicide chemistries in managing *A. tuberculatus*, novel approaches focusing on seedbank depletion are being explored, including gene drive to reduce or eliminate populations (Liu *et al.*, 2020; Schleich *et al.*, 2023; Soltani *et al.*, 2023). In general, these genetic control strategies apply more broadly to all dioecious *Amaranthus* species (Tranel & Trucco, 2009; Neve, 2018); however, utilizing such strategies requires a deeper understanding of the basis of sex determination in the species (Montgomery *et al.*, 2023).

Previous studies investigating sex determination in amaranths confirmed males were the heterogametic sex in *A. tuberculatus* (Murray, 1940; Montgomery *et al.*, 2019; Neves *et al.*, 2020), and with indistinguishable chromosomes (Grant, 1959). More recently, a draft genome of *A. tuberculatus* assembled into 841 contigs was used to develop genetic markers for genotyping sex (Montgomery *et al.*, 2019). The genome combined with K-mer counting was further used to determine male-specific contigs (∼4.6 Mb containing 147 gene models) that likely contain the sex-determining region (Neves *et al.*, 2020; Montgomery *et al.*, 2021). This draft assembly together with short-read sequencing from four dioecious amaranths was also used to identify the conservation of male-specificity of a copy of *FLOWERING LOCUS T* (Raiyemo *et al.*, 2023). Furthermore, transcriptomic analysis revealed differentially expressed genes between male and female flowers, including *MADS18*, *MADS2*, *CMB2*, *CYP710A1*, *bHLH60*, and *bHLH91* (Bobadilla *et al.*, 2023). Due to the fragmentation of the draft assembly, the orientation and order of the male-specific contigs could not be determined. Moreso, none of the genes present on the male-specific contigs were part of the genes that were differentially expressed in the transcriptomic study.

With the rapid advances in sequencing technology, chromosome-level genomic resources have been developed for three monoecious amaranths, *A. hypochondriacus* L. (Lightfoot *et al.*, 2017), *A. cruentus* L. (Ma *et al.*, 2021) and *A. tricolor* L. (Wang *et al.*, 2023), while the dioecious species have draft genomes assembled and scaffolded to pseudochromosome contiguity (Montgomery *et al.*, 2023). Using a combination of PacBio long-read sequencing, Hi-C scaffolding, Bionano optical mapping, and haplotype phasing of a male individual, we generated two high-quality chromosome-level assemblies of *A. tuberculatus*, essentially a male and female haploid assembly, both of which significantly improve on the previous draft assembly. Comparison of the two haplotype assemblies (haplomes) allowed us to identify a contiguous sex-determining region in the genome, compare structural rearrangements between the two haplomes and to chromosome-level assemblies of three monoecious amaranths, and clarify the positions of previously reported differentially expressed genes between male and female plants.

## Materials and Methods

### Plant material and growth conditions

Seeds from a herbicide susceptible accession designated “WUS” (GRIN accession number PI 698378) were sown in presoaked potting soil (Lambert LM-GPS) and watered through subirrigation. Upon reaching ∼5 cm in height, seedlings were transplanted to 16-cm pots (America Clay Works I-A650MP) filled with the same soil and grown under greenhouse conditions (25/20C and 16/8-h day/night cycles). A single flowering male plant was placed in the dark for 72 h, after which 4 g of fresh leaf tissue was sampled for PacBio HiFi library preparation and 2 g of fresh leaf tissue for Hi-C library preparation. In each case, tissue was flash frozen in liquid nitrogen and stored at -80 C until use. The same individual plant was used to harvest 4 g of fresh tissue that was immediately shipped on damp paper towels at 4 C for Bionano library preparation. Finally, a mix of four male and four female plants, assigned genotypically as described by (Montgomery *et al.*, 2021), were grown as previously described.

Root, stem, leaf, and meristem tissue was sampled from young (∼6-10 leaf) and old (flowering) plants and combined with floral tissue from all stages of development for RNA extraction (Zymo Direct-zol RNA Miniprep). Following quantification of concentration and quality, three µg of RNA was used for PacBio Iso-Seq library preparation. All samples (∼4 g) were shipped on dry ice (except for Bionano tissue, which was shipped at 4 C) to the Genome Center of Excellence at Corteva Agriscience for DNA extraction, library preparation, and sequencing.

### Genome sequencing, assembly, and annotation

The protocols for library preparation, genome sequencing, assembly, and annotation are described in detail in Supplementary Information Methods S1. The methods describing the analyses of genome characteristics including BUSCOs, transposable elements, LTR assembly index, centromeric, and telomeric repeats are provided in Methods S2.

### Genome-wide association analysis for sex in waterhemp populations

A total of 353 individuals (175 females and 178 males) derived from an artificially generated mapping population (Wu *et al.*, 2018) and previously genotyped using RAD-seq (Montgomery *et al.*, 2019) were used for GWA analysis. The single-end raw reads were demultiplexed and cleaned using the "process_radtags" command in Stacks version 2.64 (Rochette *et al.*, 2019).

Each sample obtained was aligned to Hap2 of the *A. tuberculatus* genome assembly using BWA-MEM v0.7.17 (Li, 2013). Individuals were grouped by sex, and the "gstacks" command of Stacks was used to build loci from the aligned single-end reads. We first ran the “populations” command employing a minimal filtering (--min-maf 0.05), and then assessed the level of missingness in the two data groups using VCFtools (--missing-indv) following a previously described approach (Cerca *et al.*, 2021). We filtered out samples with greater than 60% missing data from the list of individuals (popmap), and then ran the “populations” command once again on the streamlined samples utilizing additional filtering criteria (--min-maf 0.05, -p 2, -r 0.4).

The set of variants obtained were then used for downstream analyses. Principal component analysis was performed using PLINK v1.90b7 (Chang *et al.*, 2015), and GWA analysis was carried out with BLINK-C (Huang *et al.*, 2019) using the first five principal components to account for population stratification and familial relatedness. Sex of the individuals were used as binary input (phenotypes) in the analysis. Calculation of f-statistics (*FST*) was performed using VCFtools v0.1.16 (Danecek *et al.*, 2011), and linkage disequilibrium (LD) analysis was carried out using LDBlockShow v1.40 (Dong *et al.*, 2021).

### Location of previous sex markers and genes in the assemblies

A search of the previously reported primer sets (WHMS, MU-976, MU-657.2, and MU-533) that amplified a male-specific region of *A. tuberculatus* (Montgomery *et al.*, 2019, 2021) was carried out against both haplotype assemblies with BLASTN (Camacho *et al.*, 2009) using parameters: - task blastn-short -db haplotypes -query primers -outfmt 6 -out output. Reciprocal best hit (RBH) search between genes on the previously identified male-specific contigs and genes from the haplotype assemblies was carried out with MMseqs 2 (Steinegger & Söding, 2017).

### Synteny and intragenomic analysis

MCScan (Tang *et al.*, 2008) from JCVI utility libraries v1.1.22 was utilized in defining collinear gene blocks between haplotype 1 and 2 assemblies using a C-score cutoff of 0.99. MCScanX (Wang *et al.*, 2012b) was used to investigate duplicated genes in the haplotypes, and the output collinearity files were converted to “.anchors” format using “jcvi.compara.synteny” from JCVI for circos plotting. Sequence alignment between both haplotypes, and also between the haplotypes and a previous *A. tuberculatus* male draft assembly, were carried out with Minimap2 v2.24-r1122 (Li, 2021). Alignments were visualized as dotplot using D-Genies (Cabanettes & Klopp, 2018). Structural rearrangements between the two haplotypes from the previous alignment were further refined using SyRI (Goel *et al.*, 2019). Syntenic orthologues among the haplotypes and three chromosome-level monoecious *Amaranthus* species (*A. hypochondriacus*, *A. cruentus* and *A. tricolor*) were evaluated and visualized using GENESPACE v1.2.3 (Lovell *et al.*, 2022).

### Sequence divergence and detection of positive selection

We followed previously described protocols to test for the evidence of adaptive evolution for 496 single-copy orthologous genes on chromosome 1 (Jeffares *et al.*, 2014; Álvarez-Carretero *et al.*, 2023). Briefly, each orthogroup on chromosome 1 from the previous orthofinder run were aligned separately using custom perl scripts, “multiple_sequence_spliiter.pl” and “align_orthologs.pl” from Jeffares *et al.*, (2014). Synonymous (*d*S) substitution rates between the Hap1- and Hap2-linked genes (i.e., pairwise comparison using the aligned single-copy orthologous genes) were estimated with the maximum-likelihood (ML) method of Goldman & Yang (1994) implemented in the CODEML program (runmode = -2, CodonFreq = 2) in PAML package v4.10.7 (Yang, 2007). We included chromosomes 4, 7, and 16 in the analysis to evaluate synonymous divergence for autosomes. Eight genes representing outliers were each removed on chromosomes 1 and 4. We then proceeded to test for evidence of adaptive evolution for genes on chromosome 1 using a series of models. We compared the simplest or null model against a nested alternative site model (M0 vs M1a) that allows neutral sites (ω = 1) to be evaluated. We then asked if adding a third class with ω > 1 fits the data better than a model with only two classes, ω < 1 or ω = 1 (M1a vs M2a), as certain sites could be under positive selection.

Given the alignment of the single-copy genes on chromosome 1 of chromosome-level assembly of three monoecious amaranths and the two haplotypes, we used the branch models to determine if ω differs among lineages (M0 vs free-ratio), and if specific foreground branches have different ω from background branches (M0 vs two-ratio). Using the branch-site model (MAnull, ω = 1 vs MA, ω > 1), we determined if the previously defined foreground branches are more likely to contain sites under positive selection (see Fig. S14 of schematic representation of trees with defined foreground and background branches). For all hypotheses tested, we implemented the mutation-selection model (FMutSel) with observed codon frequencies used as estimates (CodonFreq = 7, estFreq = 0). The model has been indicated to account for mutation bias and selection affecting codon usage, and is preferable over other models (Yang & Nielsen, 2008; Álvarez-Carretero *et al.*, 2023). *P*-values were adjusted using the Benjamini–Hochberg method (Benjamini & Hochberg, 1995) to account for multiple comparisons.

### Expression profiling and gene ontology (GO) enrichment analysis

mRNA-sequencing data for three tissue types (mature flowers, shoot apical meristem, and floral meristems) from a previous study (Bobadilla *et al.*, 2023) were mapped to Hap2 of the *A. tuberculatus* genome using STAR aligner v2.7.10b (Dobin *et al.*, 2013). Two replicates from mature flower category were removed due to low mapping quality (26.47% and 28.44% uniquely mapped reads for replicate 2 and 4, respectively). Both samples were also removed from downstream analyses in the previous study. Gene counting was carried out using featureCounts v2.0.6 from the subread package (Liao *et al.*, 2014), and the differential expression (DE) analyses between sex for each tissue types were carried out with edgeR (Robinson *et al.*, 2009). Prior to the DE analysis, genes with zero counts were filtered (i.e., CPM values less than 1), and counts were normalized using the TMM normalization in edgeR. A negative binomial generalized log-linear model was then fitted to the normalized read counts. Genes were assessed as differentially expressed based on FDR < 0.05 and FC > 1.2 thresholds using the ‘*glmTreat*’ function within edgeR. Translated protein-coding sequences (CDS) of Hap2 were also assigned GO annotations using eggNOG-mapper v2.1.12 (Cantalapiedra *et al.*, 2021). GO term enrichment analysis was then carried out using topGO with nodeSize = 10, which is the minimal number of genes to keep per term. The enrichment test was performed using Fisher’s exact test and the “elim” algorithm. GO terms were assessed as significantly enriched at the default *p* < 0.01 threshold.

### Phylogenetic analysis of *FLOWERING LOCUS T*

A 200 bp *FT* sequence, previously reported as male-specific and conserved among three other dioecious amaranths closely related to waterhemp, was searched against the haplotype assemblies using BLAST (Camacho *et al.*, 2009). Failure to obtain perfect match to the candidate sex chromosome (Chr1) prompted further investigation. We queried the haplotype assemblies and three monoecious amaranth (*A. hypochondriacus*, *A. cruentus* and *A. tricolor*) assemblies for all homologs of *FLOWERING LOCUS T*, and also searched the 200 bp *FT* sequence against raw reads of *A. tuberculatus* from a previous study (Kreiner *et al.*, 2019) using SRA-BLAST (Leinonen *et al.*, 2011). The top hit (ranked by evalue and bitscore) and another hit were selected from each male while only the top hit for females were selected. All the *FT* sequences were aligned using MAFFT version 7 online (Katoh & Standley, 2013), and columns with less than 15% occupancy in the alignment were removed with Jalview v2.11.3.2 (Waterhouse *et al.*, 2009). A maximum likelihood tree was then constructed with the alignment in RAxML v8.2.12 (Stamatakis, 2014) using a GTRGAMMA substitution model and 1000 rapid bootstrap replicates. The resulting tree was visualized with Dendroscope v3.8.10 (Huson & Scornavacca, 2012).

### Statistical analyses

The sample sizes used for analyses are indicated in the figures. Gene count per 500 kb, TE proportion per 500 kb, and LTR insertion times data were analyzed with Kruskal-Wallis test. Post-hoc test of multiple comparisons was carried out with Conover-Iman test, following the rejection of the null hypothesis from the Kruskal-Wallis test. *P*-values were adjusted for multiple comparisons with Benjamini–Hochberg correction (Benjamini & Hochberg, 1995). The pairwise synonymous divergence (*dS*) between species (i.e., the two haplotypes and three monoecious amaranths) previously estimated with CODEML were also analyzed following the method described above, except Dunn’s test was used for multiple comparisons. All analyses were carried out with the PMCMRplus package in R v4.1.2 (R Core Team, 2021).

## Results

### Assembly metrics and genome repetitive landscape

Analysis of both haplome 1 (Hap1) and haplome 2 (Hap2) assembly completeness using BUSCO “embryophyta_odb10” database revealed 96.9% complete BUSCOs. LTR assembly index (LAI) also revealed high quality assemblies with average LAI scores for Hap1 and Hap2 at 19.19 and 20.49, respectively (Table 1). Repeat analysis using RepeatMasker revealed that 68.6% of Hap1 and 66.3% of Hap2 were made up of repetitive elements. The LTR/*Ty3* elements were the most abundant retrotransposons, representing 18.1% and 16.3% of the genome in Hap1 and Hap2, respectively (Table S1). The total repeat content and the abundance of *Ty3* elements are consistent with the 66.0% total repeats and 17.0% *Ty3* elements reported for *A. tuberculatus* draft genome assembly (Raiyemo *et al.*, 2023).

**Table 1.** Comparison of assembly statistics between the two haplomes of *A. tuberculatus* genome assemblies.

| Genome characteristics | Hap1 | Hap2 |
| --- | --- | --- |
| Assembly size (Mbp) | 613.8 | 588.8 |
| Scaffold N50 (Mbp) | 38.92 | 36.16 |
| Scaffold L50 | 7 | 7 |
| GC content (%) | 35.0 | 34.7 |
| Complete BUSCO (%) | 96.9 | 96.9 |
| Size of Ns (Mbp) | 8.10 | 9.03 |
| LTR assembly index (LAI) | 19.19 | 20.49 |
| Protein-coding genes | 23,253 | 23,160 |
| Mean gene length (bp) | 5,371 | 5,347 |
| Mean CDS length (bp) | 1,195 | 1,198 |
| Mean exon length (bp) | 284 | 285 |
| Mean exon per gene | 5.7 | 5.7 |
| Number of tRNA | 2,096 | 2,026 |
| Number of genes in orthogroups | 22,942 | 22,905 |
| Percentage (%) of genes in orthogroups | 98.7 | 98.9 |

Analysis of high-copy tandem repeats using StainedGlass heatmaps revealed that chromosomes 1, 2, 3 4, 6, and 8 appear to be submetacentric while chromosomes 9, 10, 11, 12, 14, 15, and 16 appear to be telocentric in both haplomes (Fig. S1, Fig. S2). Some regions identified as centromeric by StainedGlass were also identified as centromeric from CentroMiner (Table S2). BLAST search of the simple telomeric repeat, TTTAGGG against both haplotypes revealed telomeric repeat sequences at 29 and 28 out of the possible 32 telomeric ends for Hap1 and Hap2, respectively (Table S3). The number of *A. tuberculatus* telomeres assembled is comparable to 30 out of 34 telomeric ends reported for *A. tricolor* (Wang *et al.*, 2023).

Further annotation of the genome revealed Hap1 and Hap2 had 1,373 and 1,344 genes annotated as transcription factors (TF), respectively (Table S4 and Table S5). In addition, Hap1 had 1,346 genes annotated as disease resistance genes while Hap2 had 1,348 genes annotated as such (Table S6 and Table S7). Annotation of transfer RNAs revealed 2,096 tRNA genes in Hap1 and 2,026 tRNA genes in Hap2 (Table S8). Other non-coding RNAs in Hap1 (90 miRNAs, 890 rRNAs, 176 snRNA, 900 snoRNA) and Hap2 (93 miRNAs, 579 rRNAs, 194 snRNA, 979 snoRNA) were also annotated (Table S9).

Visualization of the genomic features analyzed above indicates an inverse relationship between gene density and LTR proportions, whereby gene-rich regions are LTR poor and vice versa (Fig. 1).

**Fig. 1.**
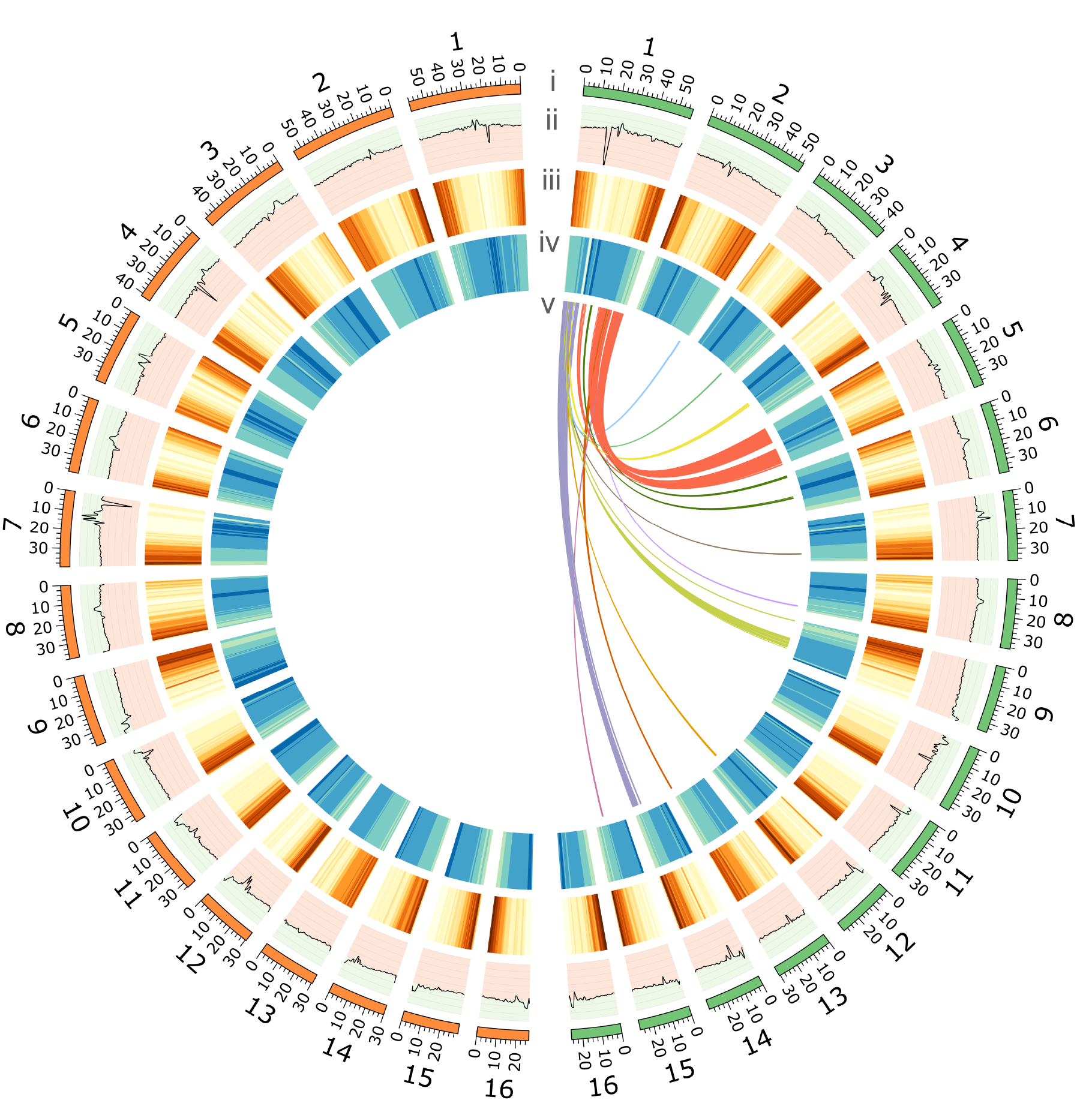
Genomic features of *A. tuberculatus* Hap1 (left) and Hap2 (right) assemblies. Circos plot depicts i) number and length (Mb) of chromosomes, ii) GC content along the chromosomes, with peaks in light green area representing GC content greater than the median (> 0.339317) and peaks in light red area representing GC content less than the median (< 0.339317), iii) gene density across the chromosomes, with brown representing gene-rich regions and yellow representing gene-poor regions, iv) LTR (long terminal repeats) density along chromosomes, with blue representing LTR-rich regions and green representing LTR-poor regions, v) inner ribbons represent duplicated genes on chromosome 1 of Hap2. Duplicated genes on chromosome 1 of Hap1 are not shown in the figure to avoid redundancy. Window size of 1 Mb and step size of 500 kb for ii-iv.

### Identification of the candidate sex-determining region

Preliminary analysis to filter out individuals with more than 60% missing data retained 44 female and 48 male samples, and the number of variants also reduced from 558,762 to 204,641.

Genome-wide association (GWA) analysis with the 92 individuals revealed that the most significant single-nucleotide polymorphisms (SNPs) associated with sex were located on chromosome 1 and spanned from 13.95 – 46.60 Mb (Fig. 2a). All identified significant SNPs resided within intergenic regions (Table S10). A QQ plot revealed no evidence of systematic bias (e.g., from analytical method, model choice, genotyping error, or population structure) in the GWA analysis (Fig. 2b). Genetic differentiation (*FST*) along chromosome 1 between females and males above the top 5% threshold spanned from 14.65 – 42.6 Mb, with a second peak at 51.6 –51.7 Mb (Fig. 2c,d). Although no clearly defined linkage blocks were observed, the sex-determining region identified above corresponds to the region indicated as containing SNPs in possible linkage disequilibrium (Fig. 2e). The region between 0 – 13.7 Mb did not show much differentiation, and thus could be considered a pseudoautosomal region (PAR) that is still actively recombining between the haplomes (Fig. 2c,d). Considering the lines of evidence above, we defined an approximate boundary of the sex-determining region (SDR) as a region of chromosome 1 on Hap2 between 13.8 – 46.6 Mb (∼ 32.8 Mb), and the Hap1 equivalent is between 14.88 – 46.0 Mb (∼ 31.12 Mb).

**Fig. 2.**
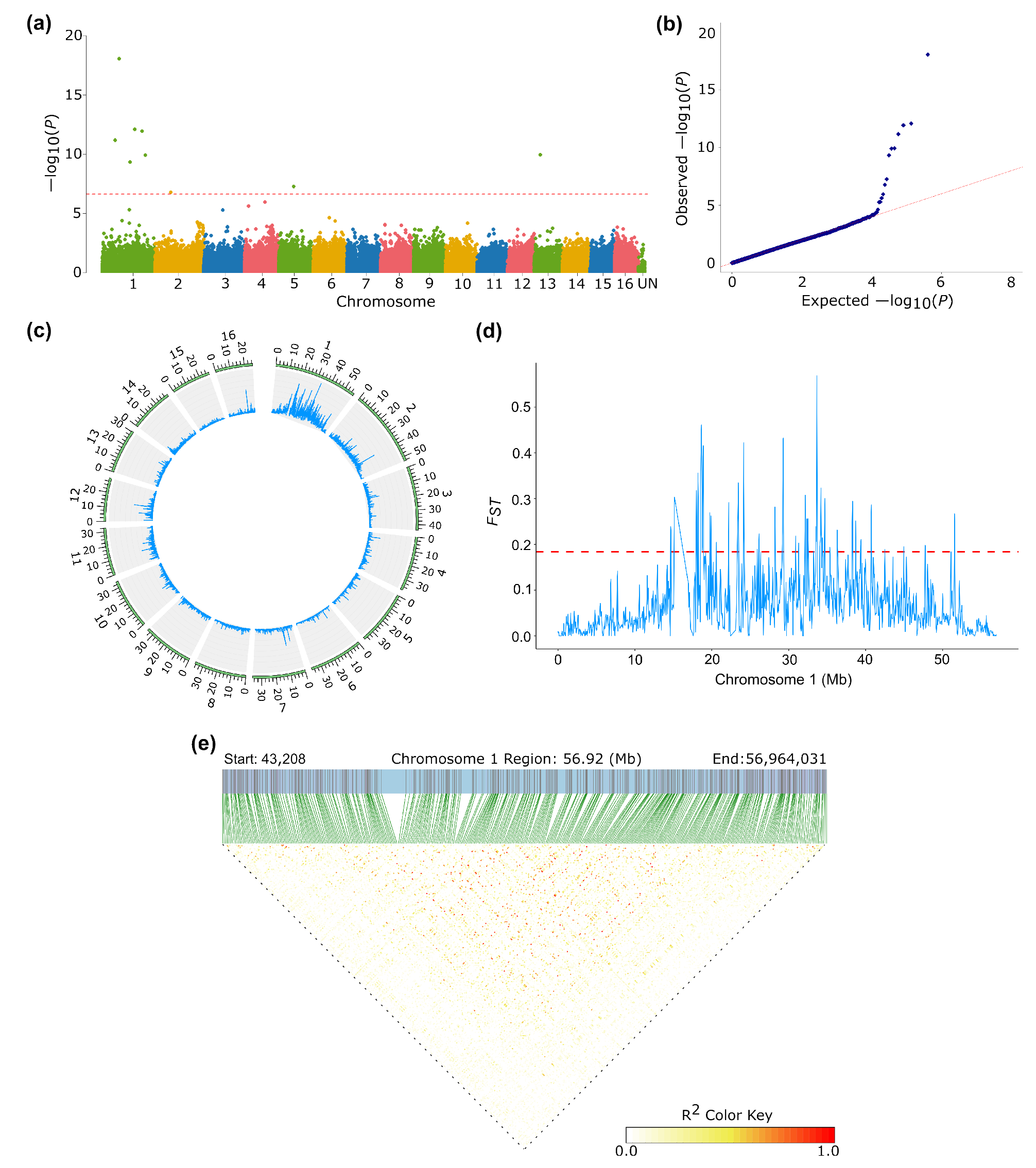
Identification of sex-determining region on chromosome 1. (a) Manhattan plot of GWA analysis using RAD-seq data from 44 females and 48 males. The dashed red line indicates Bonferroni threshold of –log10(*P*) = 6.6120. (b) Quantile-quantile (QQ) plot of the GWA analysis. (c) Fixation index (*FST)* between females and males across all 16 chromosomes (window size 100 kb; step size 50 kb). (d) *FST* between females and males for chromosome 1 with dashed red line representing the top 5% threshold at 0.1841 (window size 100 kb; step size 50 kb). (e) plot of linkage disequilibrium analysis using 375 pruned SNPs at 1 SNP/50 kb across chromosome 1.

A BLAST query of previously reported primer sets, used to amplify male-specific regions (Montgomery *et al.*, 2021), against both haplotypes also revealed perfect matches to chromosome 1 on Hap2. One set of primers (WHMS) matched to a 572 bp region (31,810,427 – 31,810,999 bp) (Fig. 3a) while primer sets MU-976, MU-657.2, and MU-533 also matched to regions within the SDR on Hap2 (Table S11). The 572 bp male-specific marker had homology to an LTR/*Ty3* retrotransposon (Fig. S3). Similarly, MU-657.2 was previously reported to have homology to a LTR/*Ty3* element from sugar beet (Montgomery *et al.*, 2019), indicating the abundance of the *Ty3* superfamily within the SDR. Sequence alignment of the previous draft *A. tuberculatus* male assembly to both haplotypes and visualization of the identified male-specific Y contigs indicated several of the contigs aligned to chromosome 1 in both haplotypes (Fig. S4 – S10). Although tig00000542 was previously proposed as one end of the male-specific Y region while tig00100752 was the other end (Montgomery *et al.*, 2021), our alignment revealed that tig00000298 and tig00000455 precedes tig00000542 while tig00100752 precedes tig00000336 and tig00000340 (Fig. S5 – S10). Structural variation between waterhemp populations could have caused these differences in the contig positions, as the previous draft genome was assembled from an individual selected from a different population than the new assembly.

**Fig. 3.**
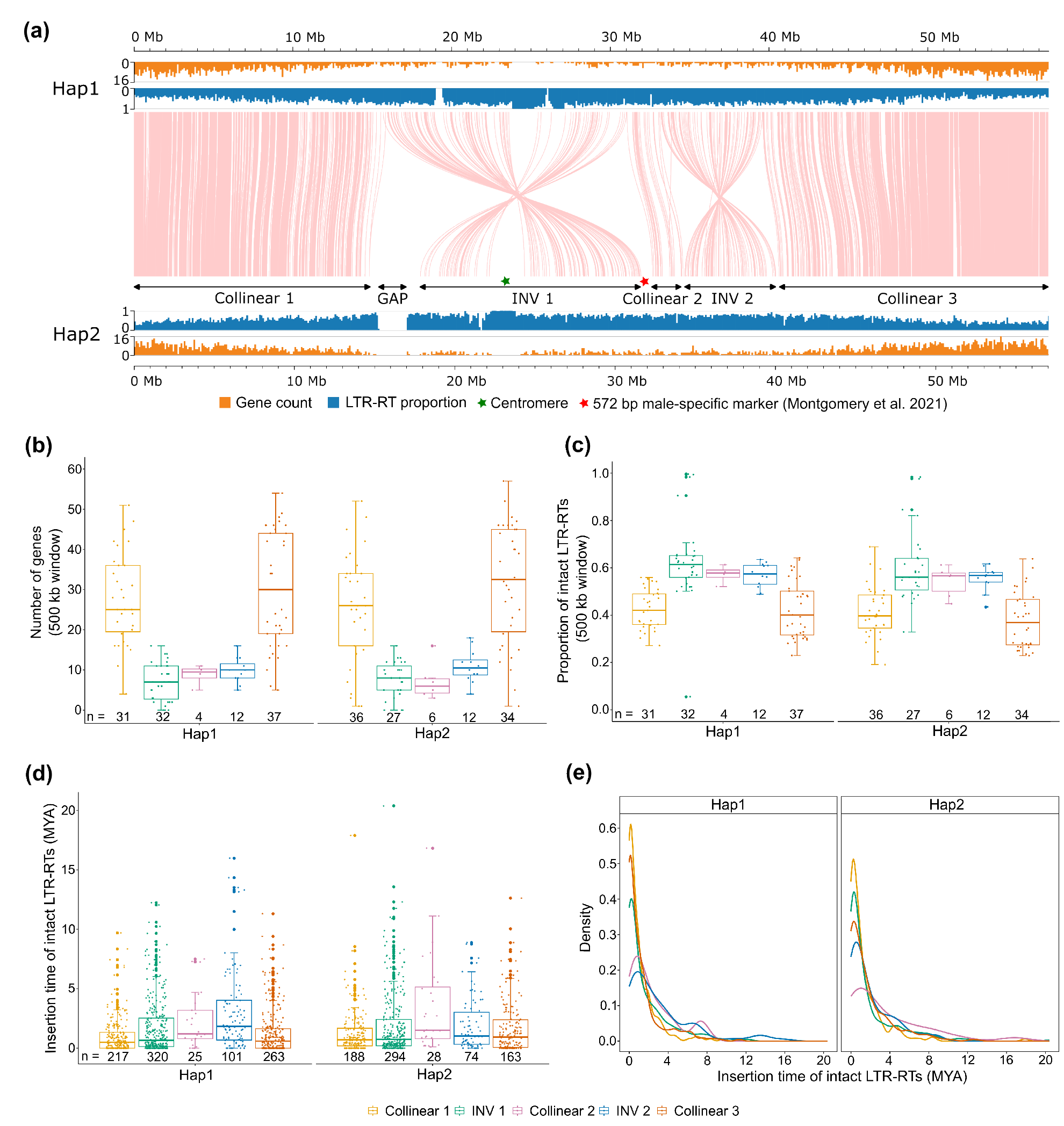
Comparative analysis between chromosome 1 of the two haplomes. (a) Synteny plot based on gene order, showing collinear regions and inversions (INV) on chromosome 1. The red asterisk represents a previously reported 572 bp male-specific marker(Montgomery *et al.*, 2021) that matched to a region on Hap2. Gene count and LTR-RT proportion were calculated based on 100 kb non-overlapping windows. (b) Number of genes across five regions (three collinear regions and two inversions) on the chromosome. Gene densities are calculated per 500 kb non-overlapping windows. (c) Proportion of intact LTR-RTs across the five regions on the chromosome. LTR-RT proportions are calculated per 500 kb non-overlapping windows. (d) Insertion time of intact LTR-RTs across the five regions. (e) Density distribution of intact LTR- RTs insertion times across the five regions on the chromosome.

There are 2,140 and 2,077 protein-coding genes on chromosome 1 of Hap1 and Hap2, of which 528 and 532 are within the SDR, respectively (Table 2). Although 5 microRNAs were identified on chromosome 1 in which three were within the putative SDR, none were haplotype-specific (Table S12). Reciprocal best hit search with the 147 genes on the previously identified male-specific contigs and the haplotype assemblies revealed 13 and 18 genes had best matches on chromosome 1 of Hap1 and Hap2 assemblies, respectively (Table S13). The genes are thus conserved between the draft assembly and the new assembly. However, 14 of the genes were syntenic between Hap1 and Hap2 based on our analysis from GENESPACE. Only 4 genes appear to be Hap2-specific with no syntenic hits to Hap1; however, these genes were annotated as proteins of unknown functions (Table S13). Taken together, the line of evidence above supports chromosome 1 as the likely sex chromosome.

**Table 2.** S. ummary statistics for chromosome 1, sex-determining region (SDR), collinear regions and inversions (INV 1 and INV 2) for the two haplotype assemblies.

|  |  | Hap1 | Hap2 |
| --- | --- | --- | --- |
| Length (bp) | Chromosome 1 | 57,604,171 | 57,035,016 |
|  | SDR | 31,116,274 | 32,798,384 |
|  | INV 1 | 15,323,107 | 13,789,911 |
|  | INV 2 | 6,357,368 | 5,640,679 |
| Number of protein-coding genes | Chromosome 1 | 2,140 | 2,077 |
|  | SDR | 528 | 531 |
|  | INV 1 | 188 | 186 |
|  | INV 2 | 114 | 111 |

### Comparative analysis between the SDR of the two haplomes

Synteny patterns between the two haplomes indicate a 1:1 relationship in gene content (Fig. S11). Out of the 20,027 gene pairs representing the reciprocal best matches between the two haplotypes, there were 1,752 pairs for chromosome 1, which were used to define the boundaries of regions within the SDR. The synteny analysis revealed two inversions on either side of a collinear region within the SDR (Fig. 3a). One inversion (INV 1) is from 15,684,914 – 31,008,021 bp on Hap1, and from 17,851,790 – 31,641,701 bp on Hap2, based on the first and last gene pairs for the inversion. The other inversion (INV 2) ranges from 33,192,221 – 39,549,589 bp on Hap1, and from 34,354,662 – 39,995,341 bp on Hap2. Further analysis of structural rearrangements using SyRI revealed 262 inversions spread across the 16 chromosomes between Hap1 and Hap2 (Table S14). INV 1 and INV 2 within the SDR were the two largest inversions out of the 262 inversions identified using SyRI (Fig. 3a).

We observed a region that was not syntenic between Hap1 and Hap2 upstream of INV 1 on Hap2 and sought to determine if the region was a translocation or duplication from elsewhere in the genome. The boundary of this region within the SDR was defined as the difference between the end coordinate of the last gene in collinear block 1 (14,660,708 bp) and the start coordinate of the first gene for INV 1 on Hap2 (17,851,789 bp), and the presence of the difference above within regions identified as “not aligned” by SyRI. Out of 14,792 regions designated as “not aligned” between the two haplomes, chromosome 1 had the highest span of a region not aligned (∼1.87 Mb; 15,186,930 – 17,059,186) (Table S17). We therefore took the “non-syntenic” region as spanning a length of 3,191,081 bp (∼3.19 Mb) on Hap2. Further analysis revealed that 1,797,397 bp (56.33%) of the region is made up of Ns and we designated the region as a “gap” (Fig. 3a).

Analysis of the genomic architecture of chromosome 1 revealed gene density for the regions (collinear 1, INV 1, collinear 2, INV 2, and collinear 3) within both Hap1 and Hap2 were statistically different with *p* < 3.35e-14 and *p* < 1.81e-10, respectively. Among pairwise comparisons of regions, only collinear 1 or 3 region had significantly higher gene densities relative to either INV 1, collinear 2, or INV 2 regions for both haplotypes (Fig. 3b, Table S16).

LTR proportions for the regions were also statistically different (*p* < 4.37e-11 and *p* < 1.60e-09, respectively). Pairwise comparison also indicated that the LTR proportions for INV 1, collinear 2, or INV 2 were significantly higher than that of collinear 1 or 3 (Fig. 3c, Table S17). Therefore, collinear 1 and 3, outside of the SDR, are gene-rich but LTR-poor while the SDR region (INV 1, collinear 2, and INV 2) is LTR-rich but gene-poor (Fig. 3b and 3c). Although analysis of insertion times revealed slight variation in the time of insertion of intact LTR retrotransposons (LTR-RTs) for collinear 2 and INV 2 relative to the other regions (Fig 3d; Table S18), both haplotypes appear to have had several of their intact LTR-RTs inserted into the regions less than 0.5 Mya, as seen in the peaks around this time (Fig 3e).

Analysis of structural rearrangements between the haplotype assemblies and three monoecious amaranths indicates a highly conserved gene order, except for a few chromosomes (Fig. 4a). Chromosome 1 in *A. tuberculatus* appears to have originated from the fusion of two chromosomes that are ancestral to chromosomes 13 and 16 from *A. hypochondriacus* and *A. cruentus*, and chromosomes 10 and 16 in *A. tricolor* (Fig. 4b and 4c). While the order of genes appears to have been maintained between monoecious species and Hap1 (Fig. 4b), the inversions discussed above appear to have occurred on Hap2 (Fig. 4c).

**Fig. 4.**
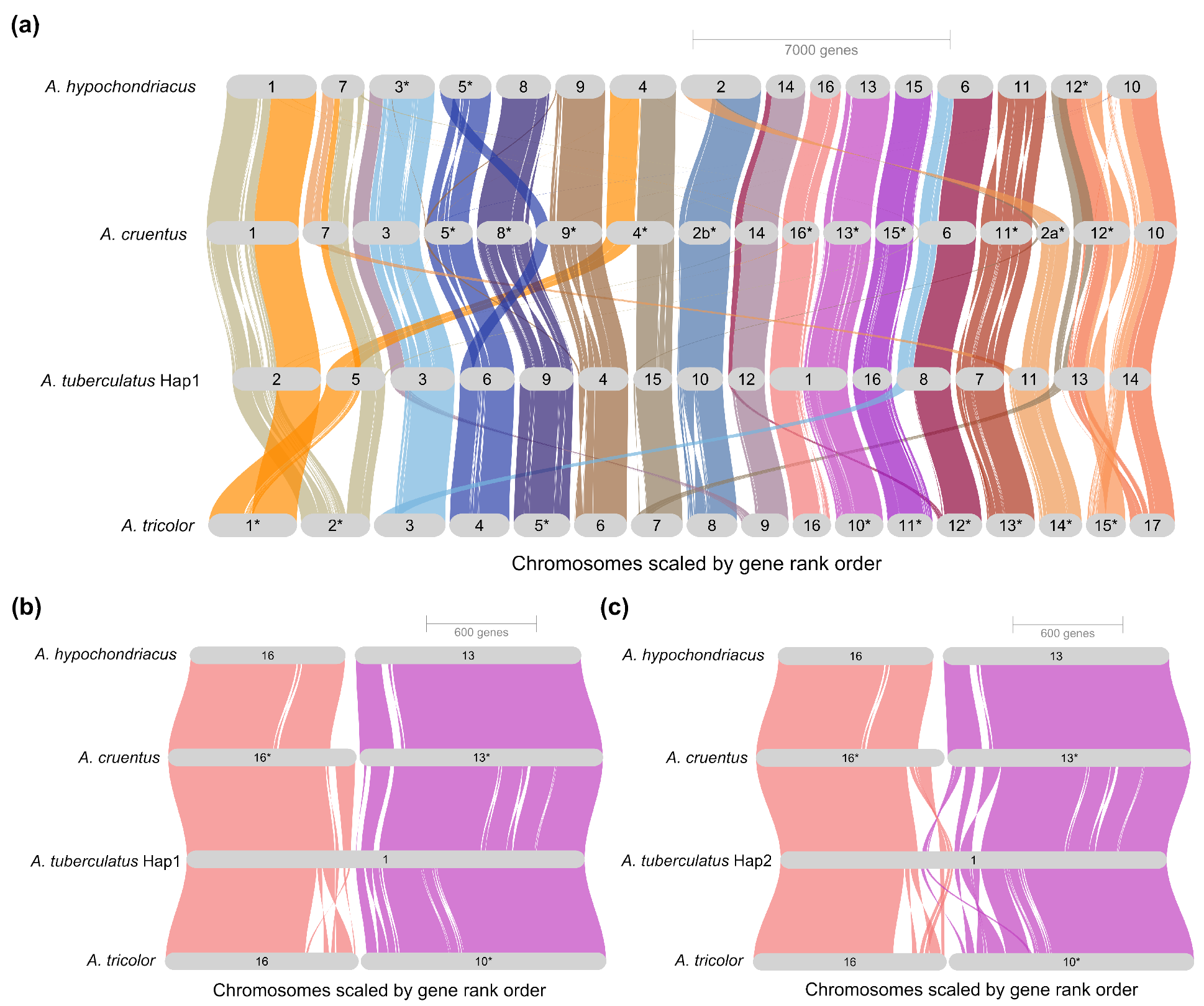
Synteny plot. (a) Synteny between the haplotype assemblies of *A. tuberculatus* and chromosome-level assemblies of three monoecious *Amaranthus* species. (b, c) Highlight of possible fusion of two separate chromosomes in *A. tuberculatus*. Asterisks indicate chromosomes that were manually inverted to keep the gene order consistent with *A. tuberculatus*.

### Sequence divergence and detection of genes under positive selection

Pairwise synonymous divergence (*ds*) was estimated between single-copy genes on chromosome 1 for the five regions (collinear 1, INV 1, collinear 2, INV 2, and collinear 3) of the haplomes and also between their orthologs in three monoecious *Amaranthus* species. The inclusion of randomly selected chromosomes 4, 7, and 16 in the analysis allowed us to assess synonymous divergence in autosomes. The mean *dS* among the chromosomes including chromosome 1 were similar, with consistency across species comparisons (Fig. S12). On chromosome 1, the mean *dS* for collinear 1, INV 1, collinear 2, INV 2, and collinear 3 for Hap1 vs. Hap2 comparison were 0.0426, 0.0283, 0.0399, 0.0308, and 0.0451, respectively. Comparisons between the five regions showed INV 1 *dS* was significantly lower than that of collinear 1 (*p* = 0.0372) and collinear 3 (*p* = 0.0142) (Table S21), indicating that recombination suppression within INV 1 occurred more recently. The higher mean *dS* for collinear 1 and 3, and the lack of statistical evidence that the two regions differ between haplotypes, indicates that the two regions are still recombining. Both regions had a comparable mean *dS* to Chr4 (0.0425), Chr7 (0.0471) and Chr16 (0.0508) (Fig. S12). For all pairwise species comparisons, collinear 1 and 3 were not statistically different in mean *dS*; however, collinear 2 and inversion 2 tend to exhibit more variation, indicating a higher sequence divergence for both regions relative to the other regions (Fig. S13). The mean *dS* of Hap1 or Hap2 to *A. tricolor* comparison was lower across the five regions, relative to the mean *dS* of Hap1 or Hap2 to *A. cruentus* or to *A. hypochondriacus* comparison, supporting previous phylogenetic evidence that indicated *A. tricolor* (subgenus *Albersia*) is more related to *A. tuberculatus* (subgenus *Acnida*) than to the other monoecious species in the subgenus *Amaranthus* (Waselkov *et al.*, 2018; Raiyemo *et al.*, 2023; Wang *et al.*, 2023).

Analysis of adaptive evolution using the M1a vs. M2a model revealed 17 genes out of 504 single-copy genes on chromosome 1 were significant (α = 0.05) for sites under positive selection (Table S20). Using the branch model, M0 vs. free-ratio, where omega (ω) was allowed to vary, there were 20 significant genes with different ω among the lineages. When Hap1 was used as the foreground branch to determine if ω differs for this branch relative to the background branches (i.e., the M0 vs. two-ratio model), there were no genes with significantly different ω.

However, when Hap2 was used as the foreground branch, only one gene encoding a serine/arginine-rich SC35-like splicing factor *SCL33* had significantly different ω, indicating it confers some fitness benefit. When both haplomes were specified as the foreground branch [i.e., (Hap1 #1, Hap2 #1)], only two genes, encoding bark storage protein B and thaumatin-like protein, had ω that differed for the specified branch. However, when the branch leading to the common ancestor of the two haplomes were included in the foreground branch (i.e., (Hap1 #1, Hap2 #1) #1)], seven genes encoding uncharacterized protein LOC110699187, bark storage protein B, LOB domain-containing protein 19-like, LRR receptor-like serine/threonine-protein kinase, thaumatin-like protein, and two copies of polygalacturonase had ω that differed for the foreground branch compared to the background branches.

Adopting the foreground and background branches used for the branch model above but utilizing a branch-site model (i.e., MAnull, ω = 1 vs. MA, ω > 1), 10 genes were more likely to contain sites with ω > 1 when Hap1 was used as the foreground branch. When Hap2 was used as the foreground branch, 16 genes were more likely to contain sites with ω > 1. When both haplomes were used as the foreground branch, 17 genes were more likely to contain sites with ω > 1. When the branch leading to the common ancestor of the two haplomes was included, 22 genes were more likely to contain sites with ω > 1 (Fig. S14, Table S20, and Table S21). Within the SDR, genes for LOB domain-containing protein 19-like, vacuolar ion transporter homolog 4- like, DEAD-box ATP-dependent RNA-helicase 39, zinc finger CCCH domain-containing protein 18-like, and prefoldin subunit 3 were more likely to be under positive selection based on the branch-site model. Taken together, our analysis above reveals genes that are potentially important in sex-specific adaptation.

### Expression analysis identifies genes involved in floral development

Clean mRNA reads (i.e., adapter trimmed and low-quality bases removed) from three tissue types (shoot apical meristem, floral meristem, and mature flower) reported by Bobadilla *et al.*, (2023) that were mapped to the Hap2 assembly had uniquely mapped reads for males ranged from 82.42 – 88.22%, and uniquely mapped reads for females ranged from 74.37 – 89.05%. Out of the 23,160 annotated Hap2 protein-coding genes, 20,259 gene were retained for DE analysis after filtering and TMM normalization (Fig. 5). Among the 1,794 genes retained on chromosome 1, two at the shoot apical meristem stage, four at the floral meristem stage, and 435 (231 upregulated and 204 downregulated) at the mature flower stage were differentially expressed between males and females (Table S22 – S24). Within the SDR on chromosome 1, two genes at the shoot apical meristem stage, three genes at the floral meristem stage, and 93 genes (55 upregulated and 38 downregulated) at the mature flower stage were differentially expressed (Table S22 – S24). Two genes within the SDR (specifically within INV 1) encoding MADS-box protein FLOWERING LOCUS C-like and LOB domain-containing protein 19-like were consistently downregulated across the three tissue types for male plants (Table S23 and Table S24).

**Fig. 5.**
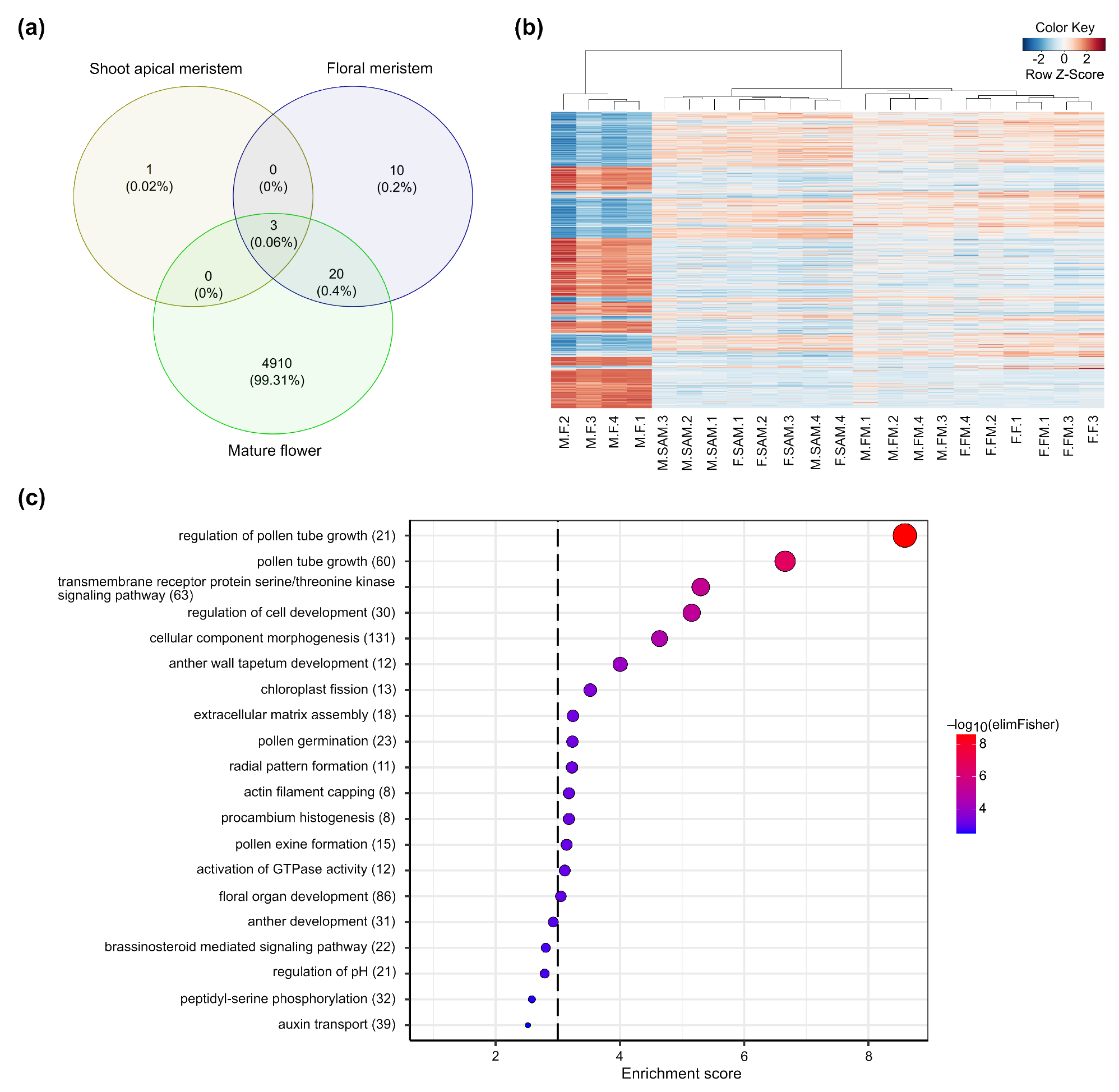
Differential gene expression analysis between male and female individuals across three tissue types. (a) Numbers of differentially expressed genes for male versus female comparison for shoot apical meristem, floral meristem, and mature flower. (b) Heatmap of log-CPM values for all 4,933 differentially expressed genes in male versus female comparison for mature flower. Samples are ordered using the hierarchical clustering method. Red depicts genes with relatively high expression, white depicts intermediate expression levels and blue represents genes with relatively low expression levels. FM: floral meristem, SAM: shoot apical meristem, M: mature flower. M or F as prefix indicate male or female, (c) Gene ontology enrichment analysis showing significantly overrepresented terms for male versus female differentially expressed genes. Values in parentheses represent the number of genes within the term. Dashed black line indicates significance level at *p*=0.001.

Gene ontology (GO) term enrichment analysis was performed to gain insight into biological processes that could be involved in sex determination. DEGs were selected based on an FDR threshold of *p* < 0.05 and FC > 1.2. Biological processes including pollen tube growth, regulation of cell development, cellular component morphogenesis, anther wall tapetum development, pollen germination, and brassinosteroid-mediated signaling pathway were identified among the top 20 enriched GO terms (Fig. 5, Table S25). *FLOWERING LOCUS C-* like and other genes including *agamous*-like *MADS-box AGL15*, *transcription factor CYCLOIDEA*-like, *transcription factor MYB44*-like, *two-component response regulator ORR24*, transcription factor *bHLH118*, probable *WRKY transcription factor 23*, and *zinc finger protein WIP2*-like were part of the “regulation of transcription factor, DNA-templated” enriched GO term (Table S26). The top five terms enriched for molecular function included transmembrane receptor protein serine/threonine kinase activity, mannan synthase activity, calmodulin binding, GTPase activator activity, and phosphatidylinositol binding (Table S27), while the top five terms enriched for cellular function included plasma membrane, apical plasma membrane, pollen tube, actin filament, and endomembrane system (Table S28).

### Male-specificity of *FLOWERING LOCUS T*

BLAST search of the 200 bp *FT* sequence, previously reported as male-specific (Raiyemo *et al.*, 2023), to the new assembly revealed no perfect matches, but 89.55% homology to a gene annotated as *HEADING DATE 3A* on chromosome 15 of both haplotypes (Hap1: 16,630,366 – 16,630,566, and Hap2: 14,349,263 – 14,349,463). Phylogenetic analysis using all homologs of *FLOWERING LOCUS T* or *HEADING DATE 3A* from the haplotype assemblies, three monoecious amaranths, and fragmented copies (< 150 bp) obtained from resequenced *A. tuberculatus* individuals via SRA-BLAST (Table S29) revealed that the copy we previously identified to be male-specific and conserved in three species closely related to *A. tuberculatus* was present in the draft assembly but absent in the new assembly (Fig. S15). A region of tig00000542 (1 – 8222 bp) where the *FT* is located within the draft assembly did not map to any region of either Hap1 or Hap2 assemblies, further indicating that the copy of the *FT* was not assembled in the new genome and could be specific to some populations.

## Discussion

We present a high-quality haplotype-resolved assembly of *A. tuberculatus*, representing the first chromosome-level assembly in the subgenus *Acnidia,* which consists of all the dioecious species in the *Amaranthus* genus. We provide evidence that chromosome 1 of the assembled genome is the sex chromosome; harboring the two largest inversions in the genome within a ∼32.8 Mb region that is gene-poor but abundant in LTR retrotransposons. Since the regions flanking the SDR are gene-rich and LTR-poor, it is possible that they are still actively recombining between the haplomes. There has been speculation over the correlation between chromosome sizes and peripheral recombination (Brazier & Glémin, 2022), with evidence in *S. latifolia* indicating that large chromosomes tend to have peripheral recombination (Yue *et al.*, 2023). Although chromosome 1 of the *A. tuberculatus* assembly is the largest chromosome (57.6 Mb), our conjecture on peripheral recombination within the flanking pseudoautosomal region is based on evidence from synteny, and on the negative relationship between gene and LTR density.

Our analysis of high-order repeats typical of centromeric regions (Melters *et al.*, 2013) indicates that chromosome 1 is either submetacentric or metacentric with the first inversion (INV 1) located in the pericentromeric region of the chromosome. Inversions have been observed within sex chromosomes of different species, including plants and animals (Wang *et al.*, 2012a; Natri *et al.*, 2019; Hearn *et al.*, 2022; Ma *et al.*, 2022), and are known evolutionary drivers of sex determination systems (Natri *et al.*, 2019). Chromosome 1 also appears to have originated from the fusion of two ancestral chromosomes following the divergence of the *Acnidia* subgenus from *Albersia*. The fusion appears to have also occurred near the centromere, perhaps as an additional mechanism to suppress recombination. Whether the chromosomal fusion event is specific to the *Acnidia* subgenus or arose independently in *A. tuberculatus* remains unclear without more chromosome-level assemblies within the *Amaranthus* genus.

The similar synonymous substitution (*dS*) rates between the haplomes for flanking regions, collinear 1 and 3 of the SDR and to autosomes, suggest that the two regions are still recombining, like the autosomal regions. Much of the variation in *dS* across species comparisons from collinear 2 and inversion 2 thus reflects the expansion of the region and accumulation of non-coding sequences due to suppressed recombination. Considering that inversion 1 has the lowest *dS* among the regions between the two haplotypes, this inversion likely occurred more recently. With no clear differences between the two haplotypes and given the similar numbers of genes present on both haplomes, *A. tuberculatus* may not have an extensive completely Y-linked region that has undergone genetic degeneration leading to loss of gene functions and deletions of genes. A similar scenario of no completely Y-linked region has been reported for spinach (Ma *et al.*, 2022). Alternatively, the Y-haplotype (i.e., Hap2) may be missing Y-specific sequences, considering the ∼1.8 Mb gap region in the assembly.

Comparison between this work and previous studies points to the influence of assembly choice on inferences drawn in sex chromosome studies. The absence of the *FT* gene fragment in this assembly that we previously reported as male-specific could either mean that it was not assembled or was specific to the population that was sequenced for the draft assembly. A search of the *FT* gene fragment in the Iso-seq data using a relaxed BLAST search parameter resulted in <90% homology to another copy. Although several authors now consider waterhemp as a single species (Waselkov & Olsen, 2014; Iamonico, 2020), it is worth noting that prior to Pratt & Clark (2001), it was considered two species, *A. tuberculatus* (primarily east of the Mississippi River) and *A. rudis* (primarily west of the Mississippi River) (Sauer, 1955, 1972) and some authors still consider them varieties, with var. *rudis* being the more weedy variety (Costea & Tardif, 2003). The population the draft assembly was derived from, designated ACR, was a weedy population from Illinois, while the population used for the new assembly, designated WUS, was from a riparian population in Ohio.

Reanalysis of the mRNA data from Bobadilla *et al.*, (2023) revealed consistent expression patterns between this work and the previous study. The two genes encoding a MADS- box protein FLOWERING LOCUS C-like and LOB domain-containing protein 19-like we reported as downregulated in males across three tissue types were also reported as downregulated in the previous study. We, however, clarify that the two downregulated genes were annotated as MADS-box transcription factor 18 (*MADS18*) and *LOB domain-containing protein 31* in that study (Fig. S16). Six genes (encoding ethylene-responsive transcription factor RAP2-7-like, CBP-diacylglycerol-glycerol-3-phosphate, protein CLT2 chloroplastic, TBC1 domain family member 8B-like, and two uncharacterized proteins) present in both the previously identified male-specific contigs of the draft assembly and within the SDR in the new assembly were also differentially expressed. Although the genes had a small difference in expression given the fold-change threshold (FC > 1.2) used in our analysis, we note that further increasing the fold-change threshold would have resulted in the same conclusion as Bobadilla et al. where none of the genes on the male-specific contigs were differentially expressed.

In sum, our study provides valuable insights into the evolution of sex chromosomes in a dioecious weedy species and adds genomic resources to the genus *Amaranthus*. We report sex-linked as well as sex-biased genes with potential roles in sex determination and flowering. Functional validation of several of the genes elicited in this study could pave the way for potential candidates for a genetic control strategy. Also, intraspecies variation in sex chromosomes or sex-determining region are being observed in some species, such as in spinach (Ma *et al.*, 2022; She *et al.*, 2023) and the human Y chromosomes (Hallast *et al.*, 2023). Future research could utilize a comparative genomics approach with other dioecious species within the genus, as well as the assembly of multiple populations of *A. tuberculatus* to further understand the variation and evolution of the sex-determining region.

## Acknowledgements

This work was supported by the International Weed Genomics Consortium with funding from Foundation for Food & Agriculture Research (FFAR grant number DSnew-0000000024), Bayer Crop Science, Corteva Agriscience, Syngenta, BASF, and the Global Herbicide Resistance Action Committee. We also acknowledge funding from the USDA National Institute of Food and Agriculture (Grant number 2022-67013-36142 to PJT).

## Author contributions

P.J.T., T.A.G., and E.L.P. conceived the research study. J.S.M. and S.M. grew plants, harvested, and prepared samples for shipping to the sequencing center. V.L. and K.F. performed DNA and RNA extraction, PacBio HiFi, Bionano DLS, Hi-C Seq, Iso-Seq sequencing, as well as genome data integration and assembly. E.L.P. and L.C. annotated the genome and provided genome assembly metrics. D.A.R. carried out genome-wide association analysis, adaptive evolution analysis, transposable elements analysis, comparative genomics analyses and statistical analyses with supervision of P.J.T. D.A.R. wrote the manuscript. All authors read, revised, and approved the manuscript.

## Data availability

Data that supports the findings of this study are available as supplementary materials. Sequencing data analyzed in this study are available through the National Center for Biotechnology Information (NCBI) (PRJNA1085752). Genome resources including FASTA files and accompanying GFF annotation files have been deposited on NCBI (PRJNA1089550), CoGe (67784 and 67785), and the International Weed Genomics Consortium online database WeedPedia (https://www.weedgenomics.org/species/amaranthus-tuberculatus/).

