## Supplementary Figure 1 - 16 for "A phased chromosome-level genome assembly provides insights into the evolution of sex chromosomes in *Amaranthus tuberculatus*"

The following Supporting Information is available for this article:

**Fig. S1** Heatmaps of tandem repeat structures for each of the 16 chromosome of haplome 1 assembly. Higher-order repeats represented by brightly-colored region indicates the centromere.

**Fig. S2** Heatmaps of tandem repeat structures for each of the 16 chromosome of haplome 2 assembly. Higher-order repeats represented by brightly-colored region indicates the centromere.

**Fig. S3** Search of the 572 bp male-specific marker to a database of transposable element encoded proteins. The marker matched to two hits with about 79.54% or 94.23% of the sequences masked. Softmasked sequences are represented with lowercase.

**Fig. S4** Dotplot of the alignment between 23 contigs in Montgomery et al. (2021) and the two haplotype assemblies.

**Fig. S5** Dotplot of alignment between tig00000298 (425,909 bp) and the two haplotype assemblies.

**Fig. S6** Dotplot of alignment between tig00000336 (82,146 bp) and the two haplotype assemblies.

**Fig. S7** Dotplot of alignment between tig00000340 (57,989 bp) and the two haplotype assemblies.

**Fig. S8** Dotplot of alignment between tig00000455 (808,180 bp) and the two haplotype assemblies.

**Fig. S9** Dotplot of alignment between tig00000542 (659,802 bp) and the two haplotype assemblies.

**Fig. S10** Dotplot of alignment between tig00100752 (1,504,067 bp) and the two haplotype assemblies.

**Fig. S11** Genomic features of the two haplotype assemblies of *A. tuberculatus*. (a) Hi-C contact map haplome 2 (alternative assembly). (b) Hi-C contact map of haplome 1 (reference assembly). (c) Dotplot of base alignment between the two haplomes. (d) synteny pattern between both haplomes indicating a 1:1 relationship in gene content.

**Fig. S12** Pairwise synonymous divergence ($d_s$) between single-copy genes on chromosomes 1, 4, 7, and 16 of the haplotype assemblies and their orthologs in three monoecious *Amaranthus* species. The number of genes for each chromosome: Chr1 = 496, Chr4 = 365, Chr7 = 341, Chr16 = 237.

**Fig. S13** Pairwise synonymous divergence ($d_s$) between single-copy genes on chromosome 1 of the haplotype assemblies and their orthologs in three monoecious *Amaranthus* species. The number of genes for each region: collinear 1 = 183, inversion 1 = 30, collinear 2 = 5, inversion 2 = 20, collinear 3 = 258.

**Fig. S14** Schematic representation of trees displaying branches used as foreground (red color) in CODEML analysis. (a) Haplotype 1 was used as the foreground branch while others were background branches ((tuberculatus1 #1,tuberculatus2),(cruentus,hypochondriacus),tricolor); (b) Haplotype 2 was used as the foreground branch while others were the background branches ((tuberculatus1,tuberculatus2 #1),(cruentus,hypochondriacus),tricolor); (c) Both haplotypes were used as the foreground branches ((tuberculatus1 #1,tuberculatus2 #1),(cruentus,hypochondriacus),tricolor); (d) Both haplotypes including the branch leading to their common ancestor were used as foreground branches ((tuberculatus1 #1,tuberculatus2 #1) #1,(cruentus,hypochondriacus),tricolor);.

**Fig. S15** Phylogenetic tree of *FLOWERING LOCUS T* copies from the haplotype assemblies and three monoecious species (*A. hypochondriacus* - AH, *A. cruentus* – Amacr and *A. tricolor* – Amatr). Numbers above branches indicate bootstrap support values.

**Fig. S16** Sequence alignment of MADS-box transcription factor 18 (MADS18) and MADS-box protein FLOWERING LOCUS C-like (FLC-like), and phylogenetic tree of LOB domain-containing protein (LBD). (a) Sequence alignment between MADS18 in Bobadilla *et al.*, (2023) and FLC-like in this work. The table below indicates FLC-like annotations from different databases (b) Phylogenetic tree of LBD. The tree leaves represent the various copies of LBD including copies from Bobadilla *et al.*, (2023). Leaves in red represent copies downregulated in males in that study and this work. Numbers above branches indicate bootstrap support values.

**Methods S1** Library preparation, sequencing, assembly, and annotation.

**Methods S2** Genome characteristics and repeat analysis.

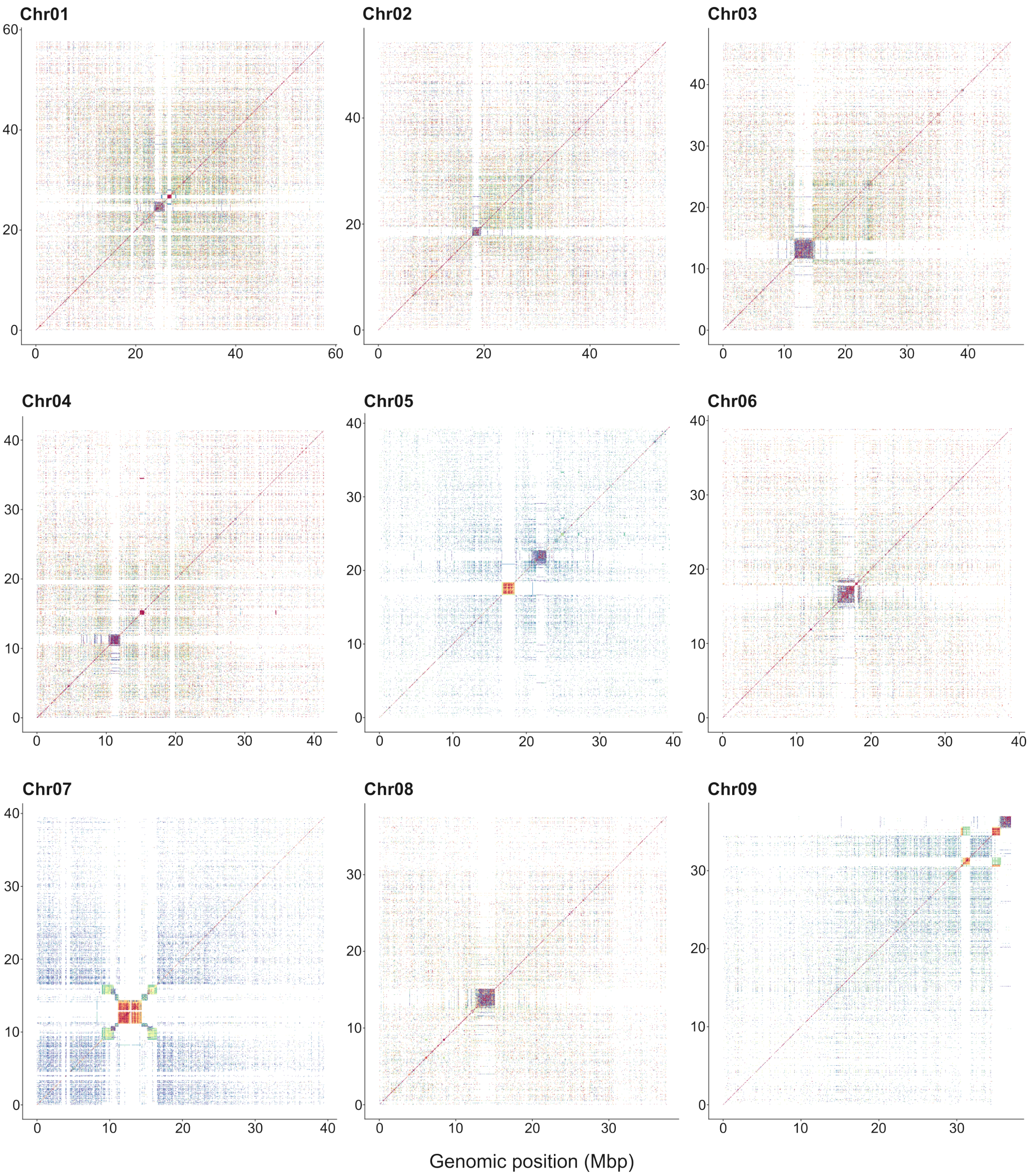
Chr01
Chr02
Chr03
Chr04
Chr05
Chr06
Chr07
Chr08
Chr09
Genomic position (Mbp)

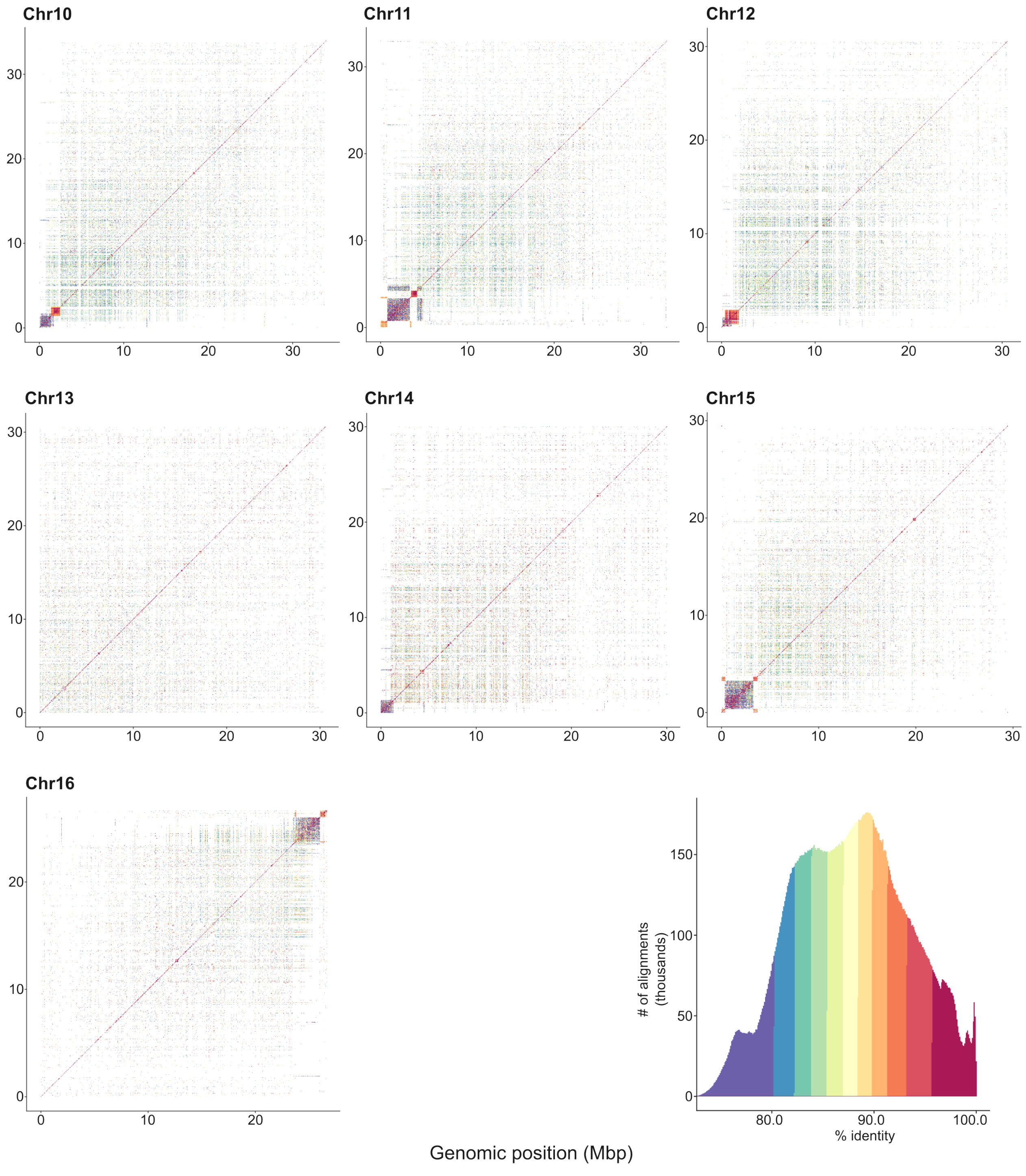


**Fig. S1** Heatmaps of tandem repeat structures for each of the 16 chromosome of haplome 1 assembly. Higher-order repeats represented by brightly-colored region indicates the centromere.

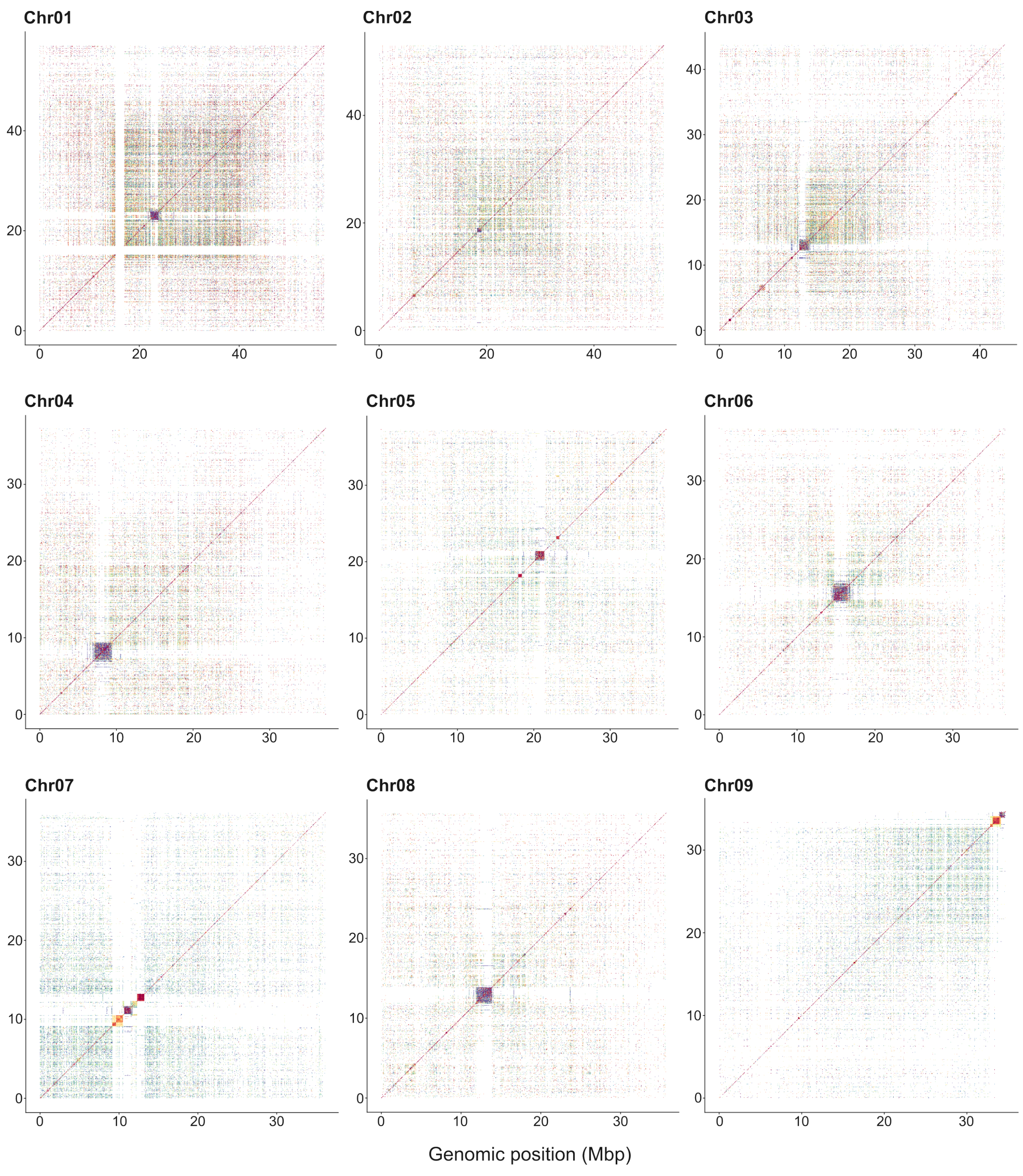
Chr01
Chr02
Chr03
Chr04
Chr05
Chr06
Chr07
Chr08
Chr09
Genomic position (Mbp)

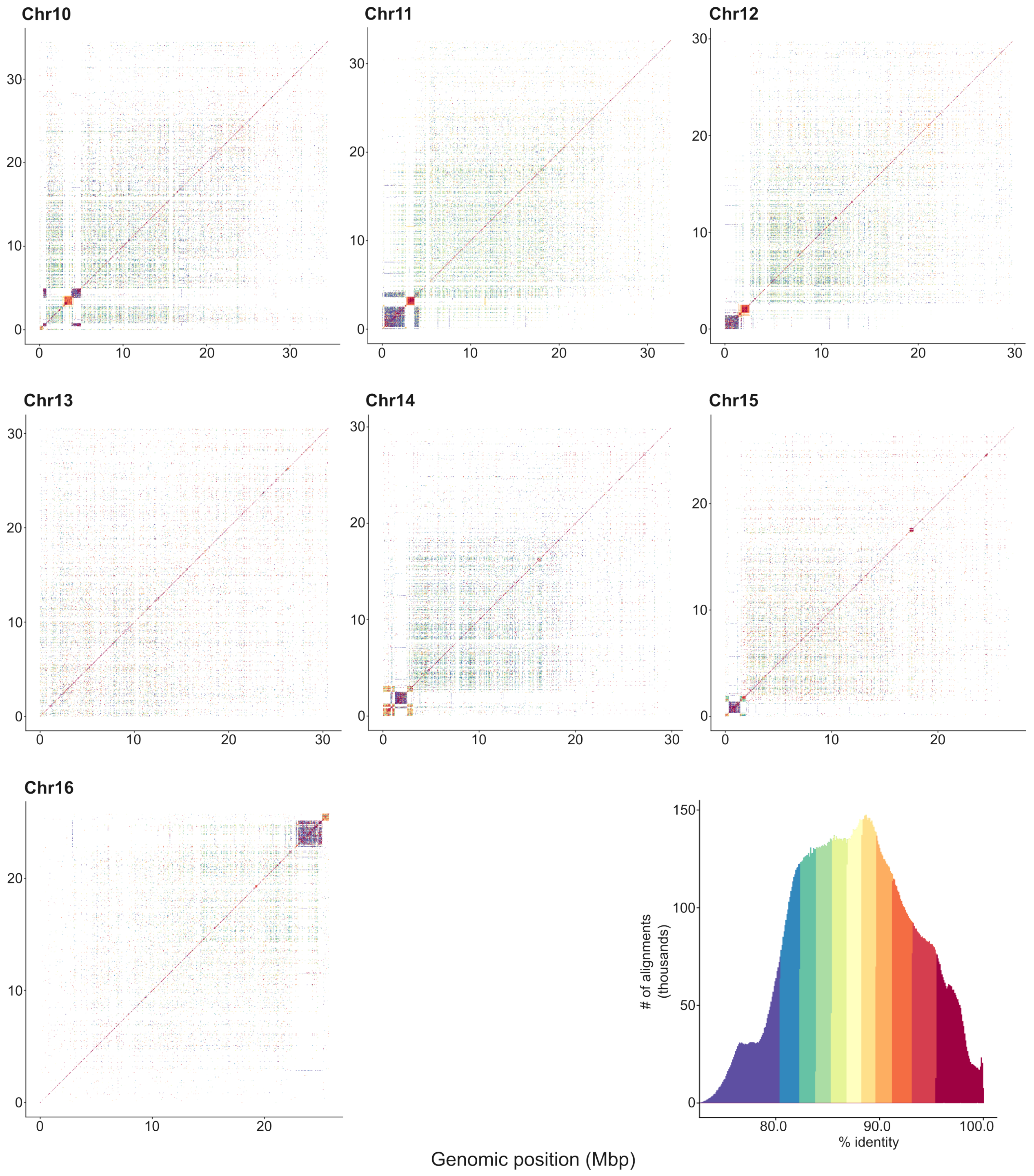


**Fig. S2** Heatmaps of tandem repeat structures for each of the 16 chromosome of haplome 2 assembly. Higher-order repeats represented by brightly-colored region indicates the centromere.

```
Masking Simple Repeats...
   - Tandem Repeats: 0
Masking Repeat Proteins...
   - Protein Hits = 2
Done!
5.57user 1.12system 0:06.76elapsed 99%CPU (0avgtext+0avgdata 799780maxresident)k
0inputs+176outputs (0major+745896minor)pagefaults 0swaps
```

Masked Sequence

| pValue | Score | Method | SeqID | Begin | End | | Repeat | Type | Begin | End |
|---|---|---|---|---|---|---|---|---|---|---|
| 0.00e+00 | 295 | WUBlastX | WHMS | 6 | 461 | - | RETROSAT4_gagpo | LTR/Gypsy | 1090 | 1241 |
| 0.00e+00 | 253 | WUBlastX | WHMS | 6 | 545 | - | SZ-63_pol | LTR/Gypsy | 685 | 871 |

```
>WHMS_572bp_marker
ATGACatcattccctcactgagtcctaagtagtcaaaccacaaaatccatagagatcgaa
tcccacttccaatcagggacaagcaagggttgaagcagtcctgaactctttttcctttttg
aattttacctgctgacataataaacacttagacacaaactctgatacatcttttcttagc
cctttccaccagattagctctctaatttcttcaatcattttatcacgaccaggatgtaaa
tgaaacaagctttgagtacctactttcaaaatgtcattcttcaactcgagttgatttgga
acgcacaacctaccattcatccttagctcccctctctcccccaactcaaaagcattaatc
ttattagcttcatctctatgtatcaattccacaacctctagatcctcaagttgtgcctca
atgatatgatcaaacaaggacagttgcacagtcattgctcccaaggagtgaccaacctct
ggccgtgtaaaatctctaacccaaagttctgcacctcacgatacaactcccatggtacag
tcatcAAAGCATTTAGACTCACTCTTGGTCTCC
```

**Fig. S3** Search of the 572 bp male-specific marker to a database of transposable element encoded proteins. The marker matched to two hits with about 79.54% or 94.23% of the sequences masked. Softmasked sequences are represented with lowercase.

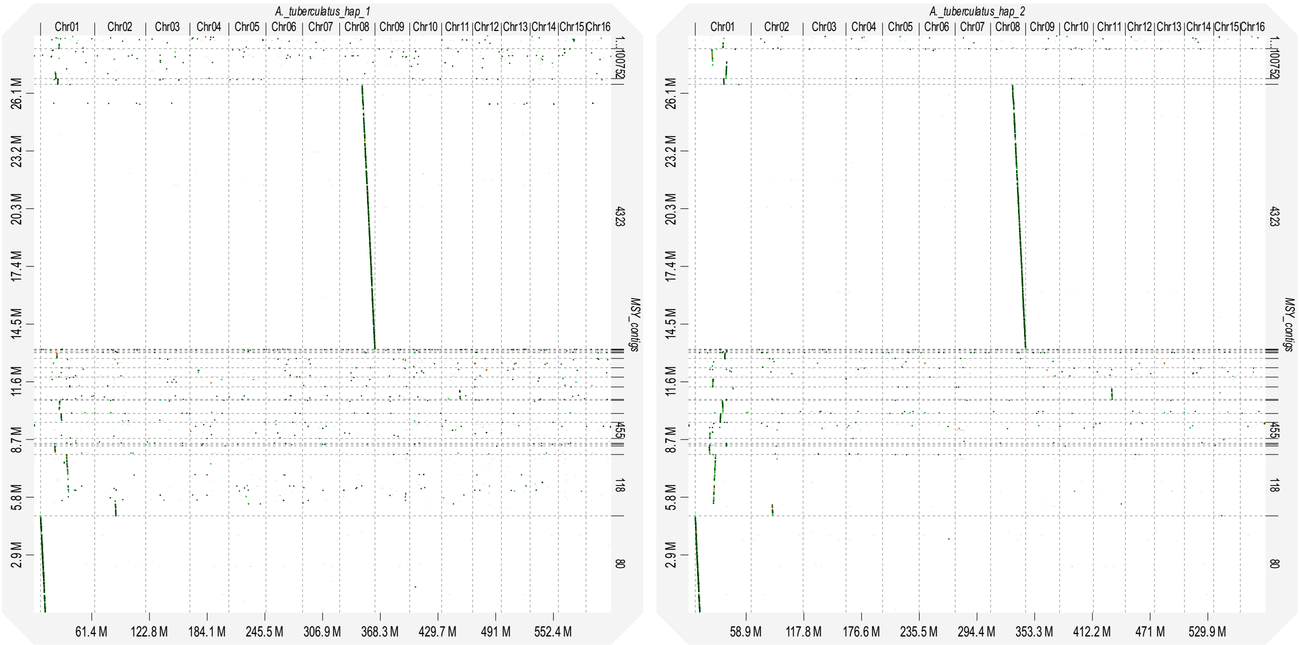


**Fig. S4** Dotplot of the alignment between 23 contigs in Montgomery et al. (2021) and the two haplotype assemblies.

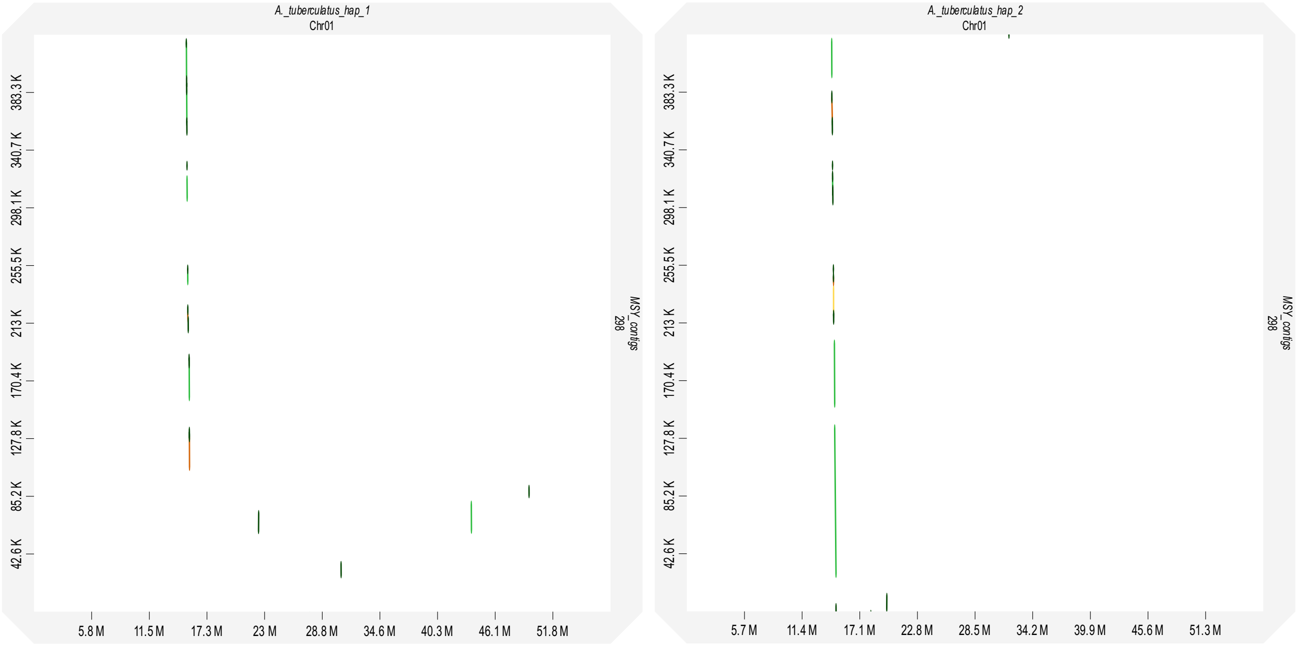


**Fig. S5** Dotplot of alignment between tig00000298 (425,909 bp) and the two haplotype assemblies.

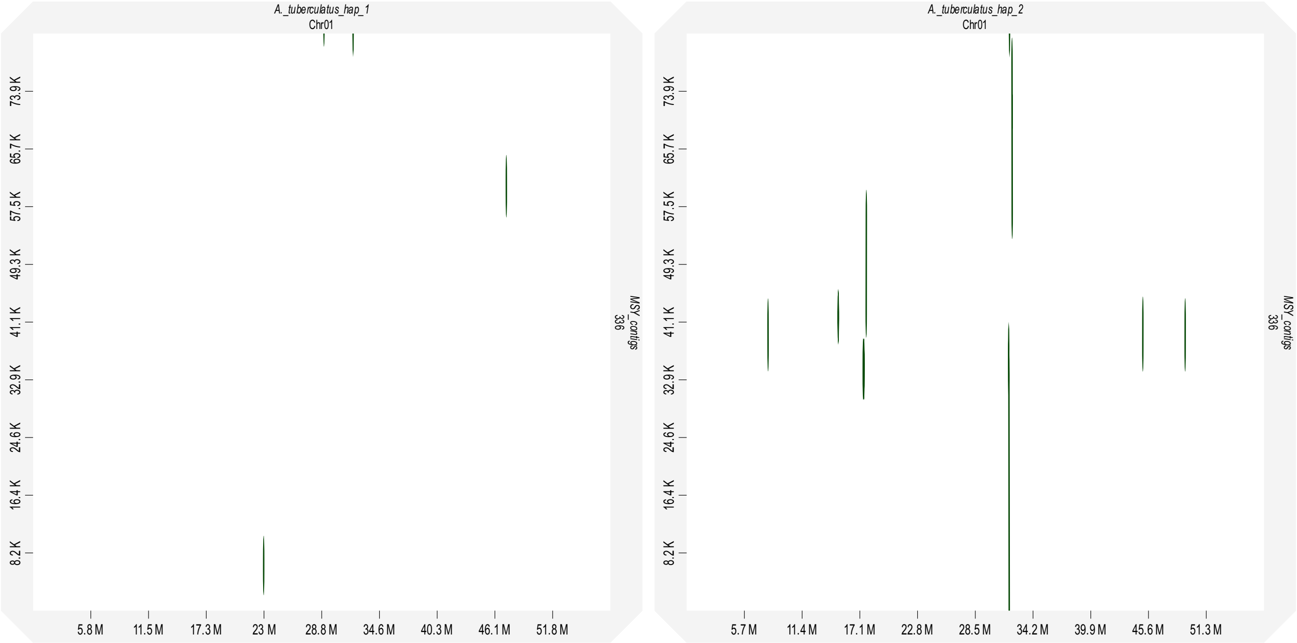


**Fig. S6** Dotplot of alignment between tig00000336 (82,146 bp) and the two haplotype assemblies.

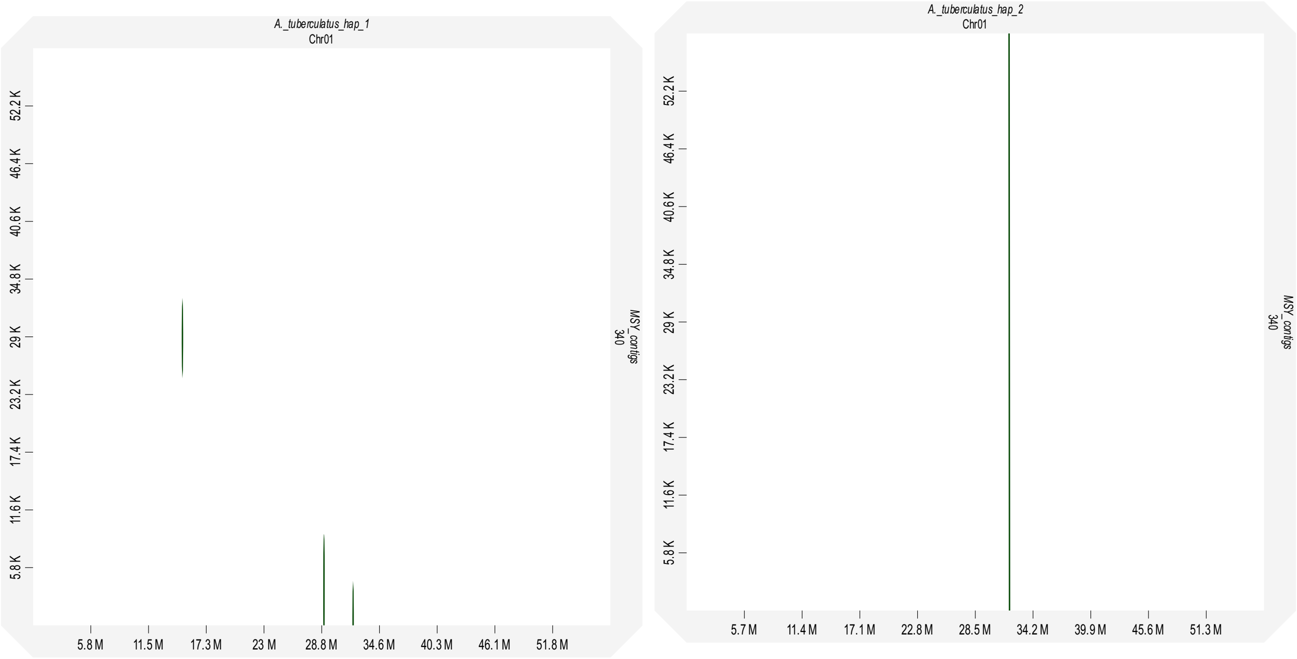


**Fig. S7** Dotplot of alignment between tig00000340 (57,989 bp) and the two haplotype assemblies.

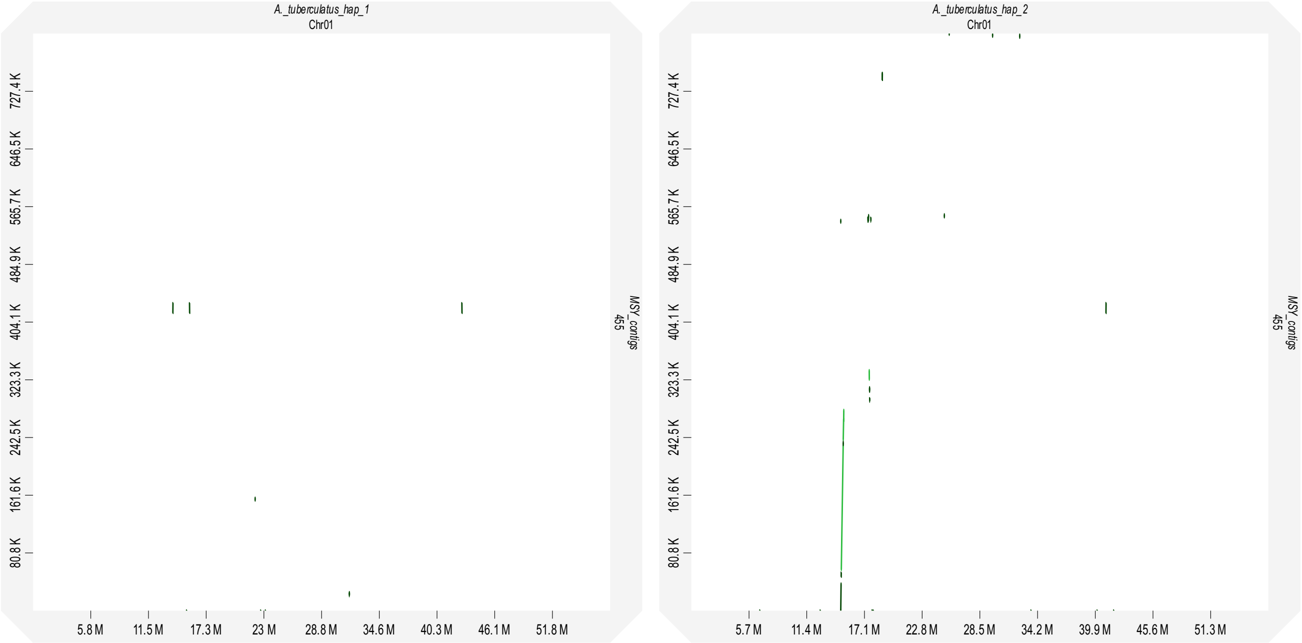


**Fig. S8** Dotplot of alignment between tig00000455 (808,180 bp) and the two haplotype assemblies.

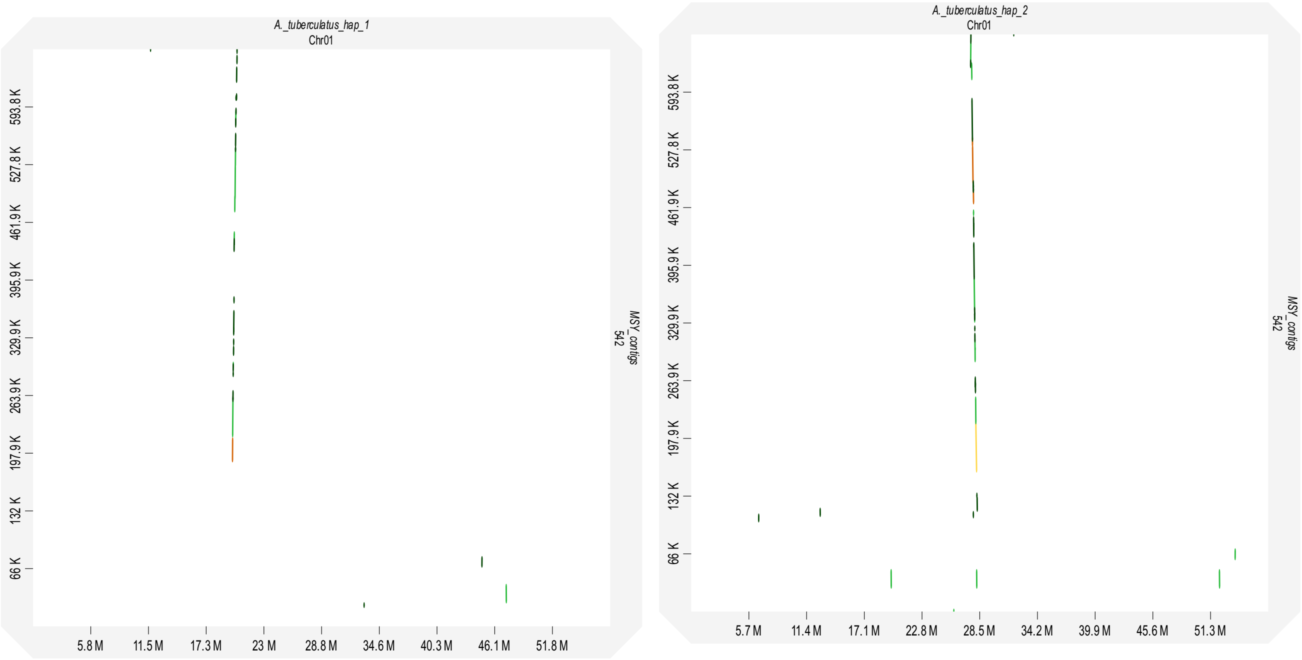


**Fig. S9** Dotplot of alignment between tig00000542 (659,802 bp) and the two haplotype assemblies.

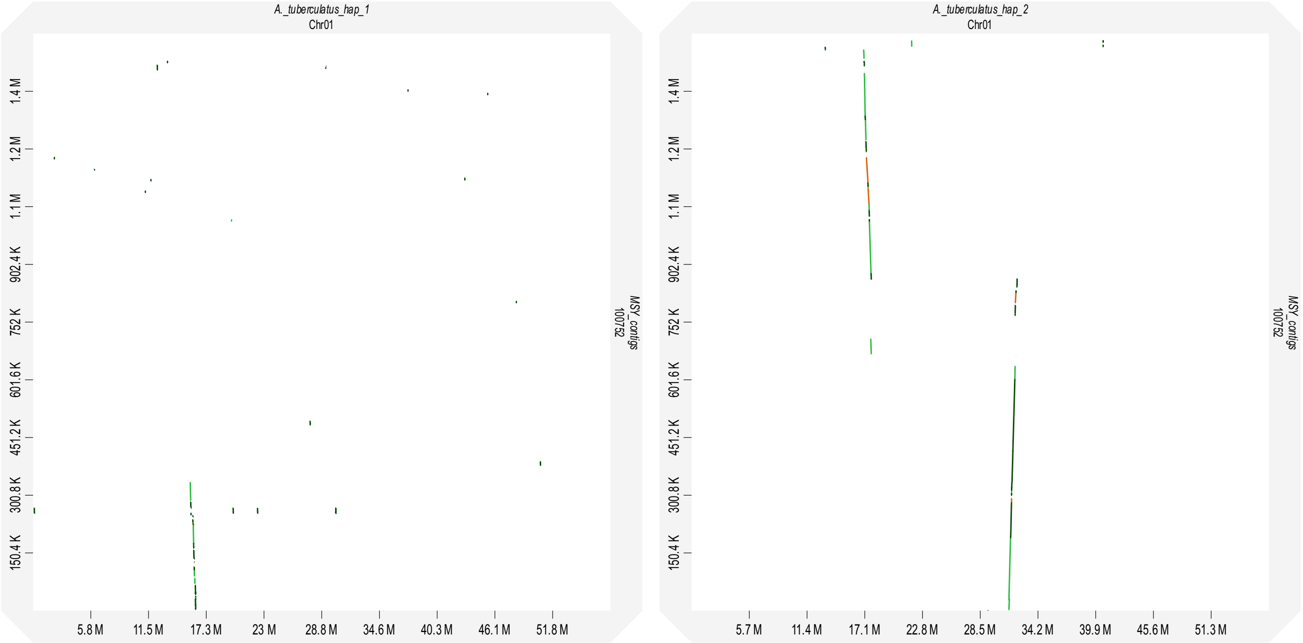


**Fig. S10** Dotplot of alignment between tig00100752 (1,504,067 bp) and the two haplotype assemblies.

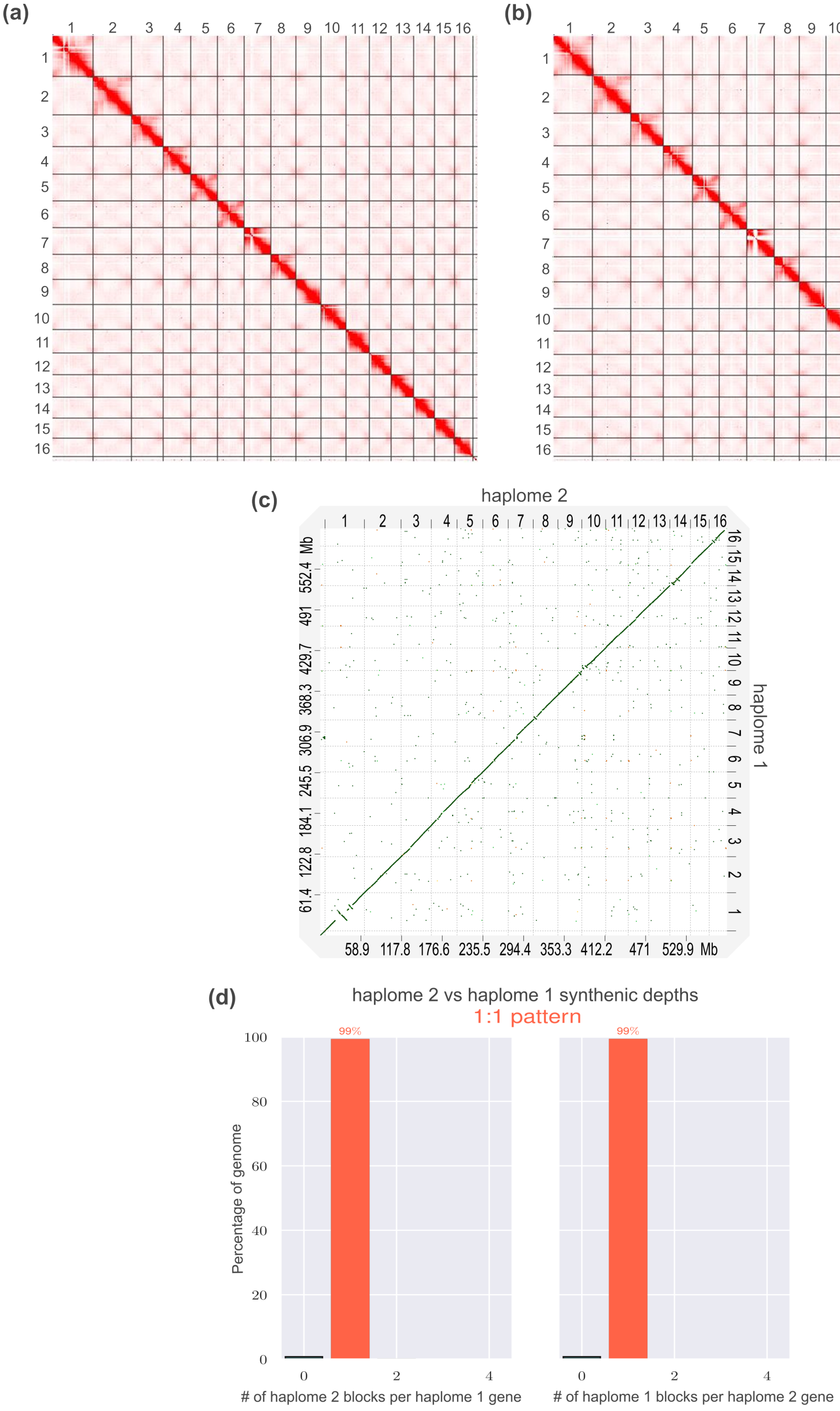


**Fig. S11** Genomic features of the two haplotype assemblies of *A. tuberculatus*. (a) Hi-C contact map haplome 2 (alternative assembly). (b) Hi-C contact map of haplome 1 (reference assembly). (c) Dotplot of base alignment between the two haplomes. (d) synteny pattern between both haplomes indicating a 1:1 relationship in gene content.

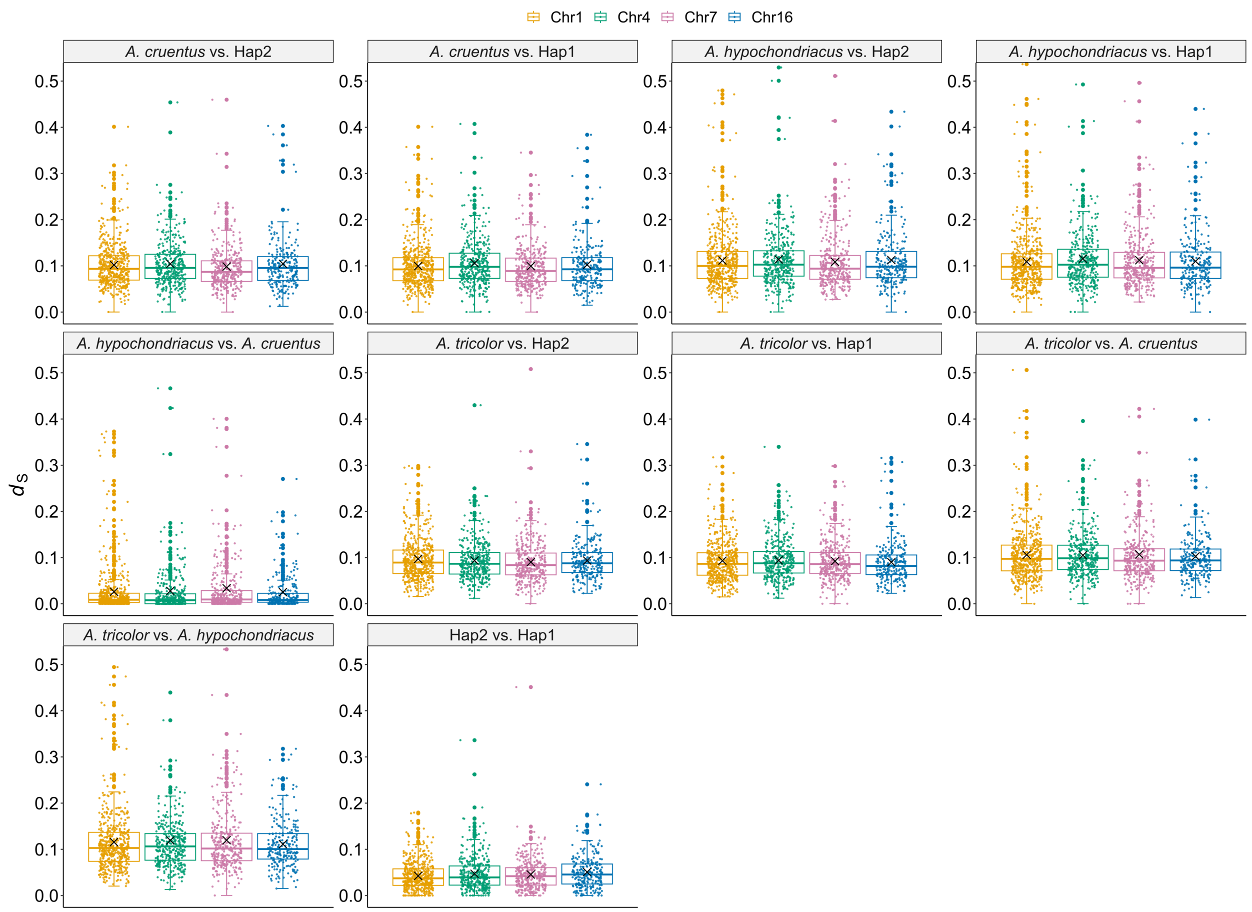


**Fig. S12** Pairwise synonymous divergence ($d_s$) between single-copy genes on chromosomes 1, 4, 7, and 16 of the haplotype assemblies and their orthologs in three monoecious *Amaranthus* species. The number of genes for each chromosome: Chr1 = 496, Chr4 = 365, Chr7 = 341, Chr16 = 237.

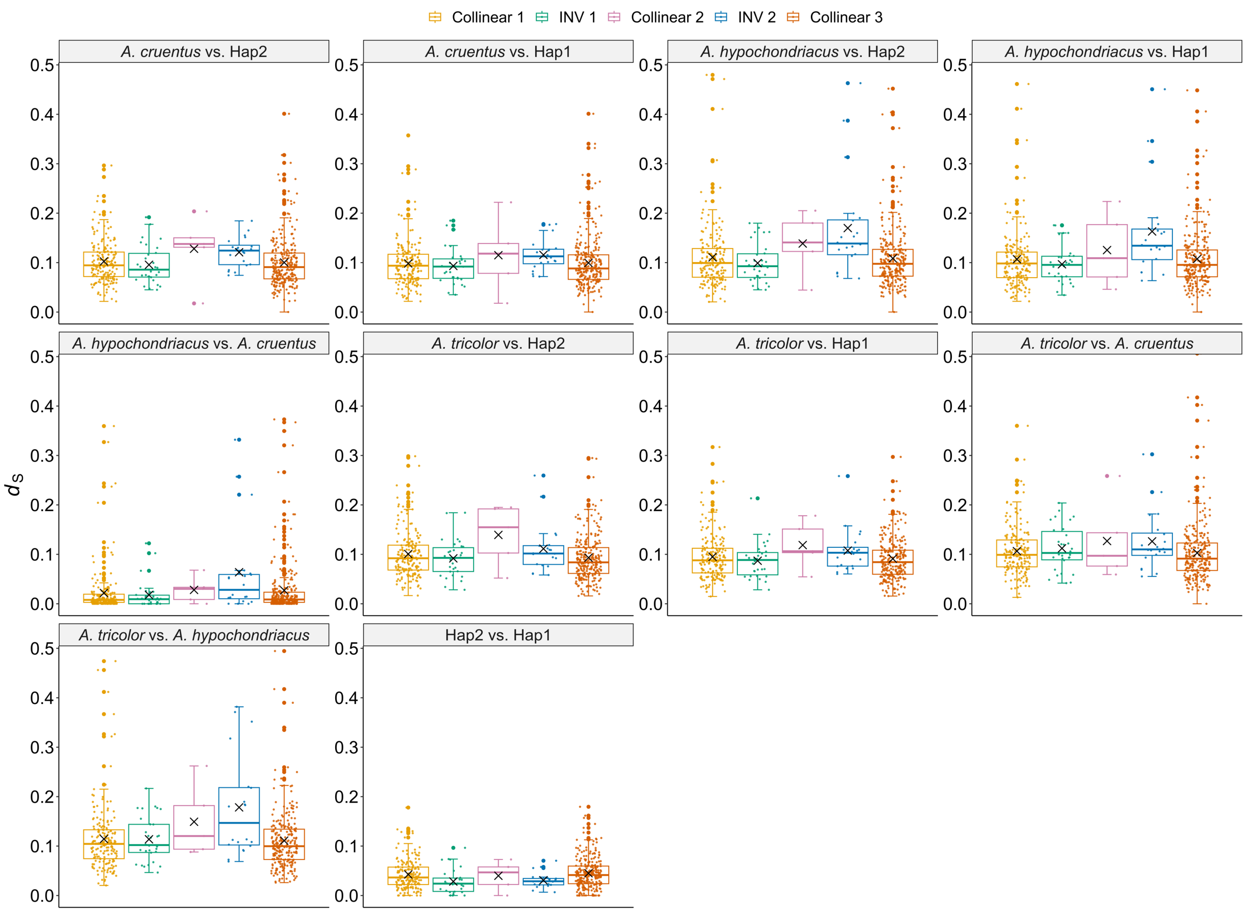


**Fig. S13** Pairwise synonymous divergence ($d_s$) between single-copy genes on chromosome 1 of the haplotype assemblies and their orthologs in three monoecious *Amaranthus* species. The number of genes for each region: collinear 1 = 183, inversion 1 = 30, collinear 2 = 5, inversion 2 = 20, collinear 3 = 258.

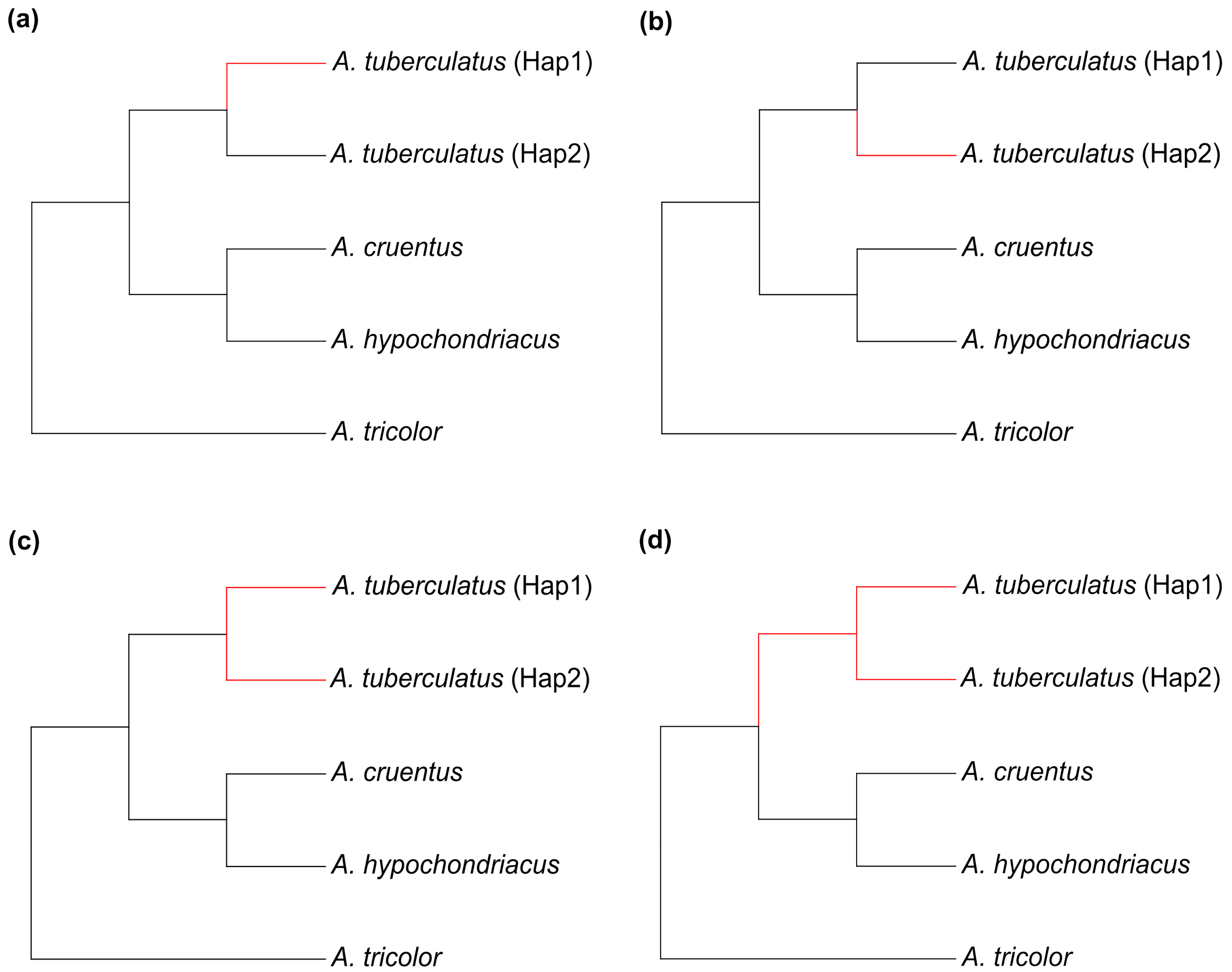


**Fig. S14** Schematic representation of trees displaying branches used as foreground (red color) in CODEML analysis. (a) Haplotype 1 was used as the foreground branch while others were background branches ((tuberculatus1 #1,tuberculatus2),(cruentus,hypochondriacus),tricolor);. (b) Haplotype 2 was used as the foreground branch while others were the background branches ((tuberculatus1,tuberculatus2 #1),(cruentus,hypochondriacus),tricolor);. (c) Both haplotypes were used as the foreground branches ((tuberculatus1 #1,tuberculatus2 #1),(cruentus,hypochondriacus),tricolor); (d) Both haplotypes including the branch leading to their common ancestor were used as foreground branches ((tuberculatus1 #1,tuberculatus2 #1) #1,(cruentus,hypochondriacus),tricolor);.

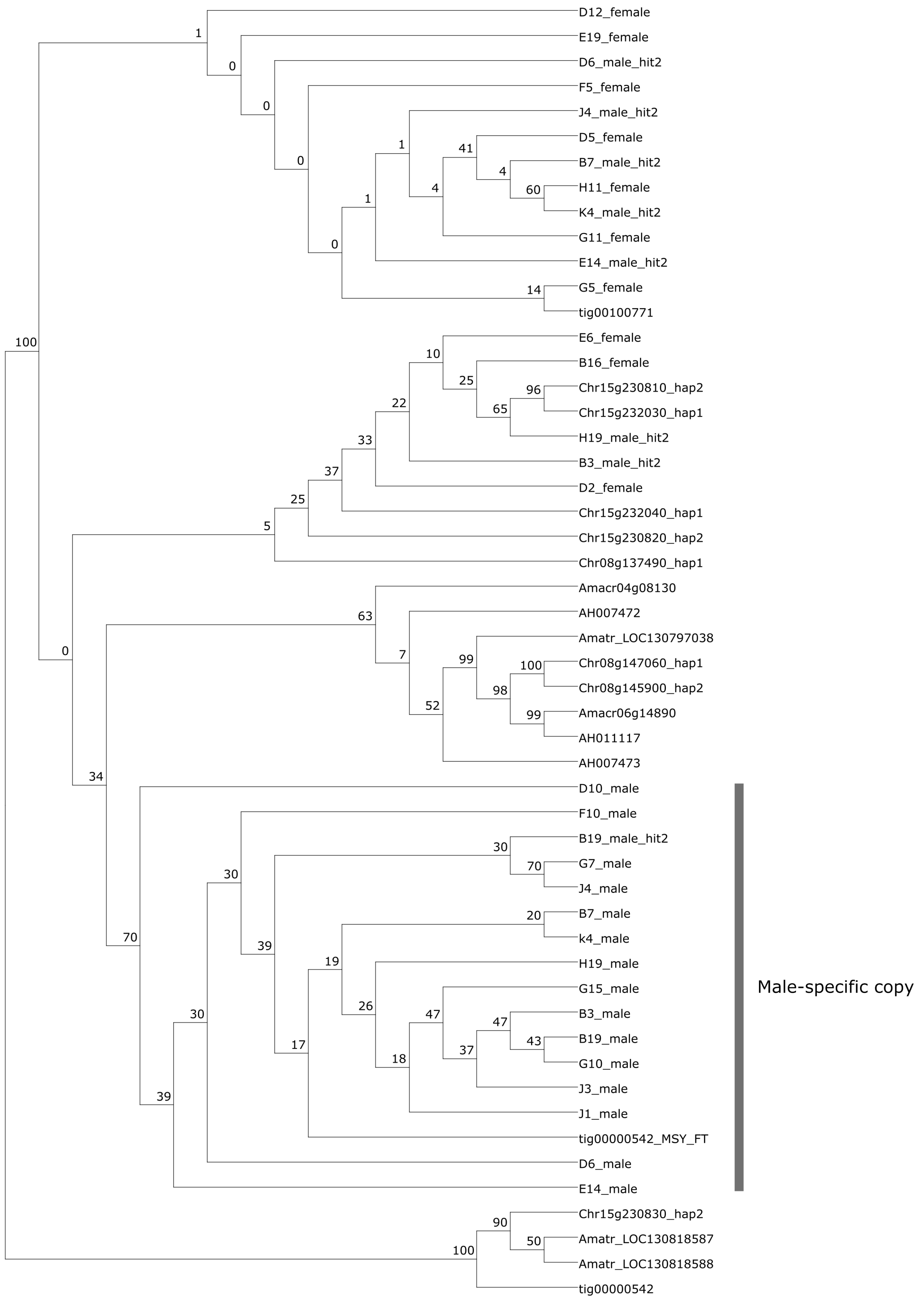


**Fig. 15** Phylogenetic tree of *FLOWERING LOCUS T* copies from the haplotype assemblies and three monoecious species (*A. hypochondriacus* - AH, *A. cruentus* – Amacr and *A. tricolor* – Amatr). Numbers above branches indicate bootstrap support values.

(a)

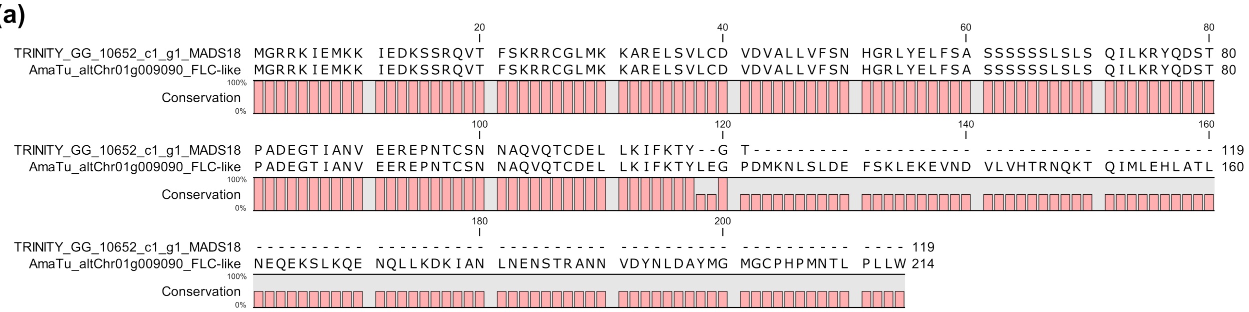


| Database | Organism | Top hit by evalue/bitscore (% identity) |
|---|---|---|
| STRING v12.0 | *Arabidopsis thaliana* | AGL27 (39.9%) |
| | *Arabidopsis thaliana* | FLC (45.0%) |
| | *Arabidopsis thaliana* | AP1 (35.3%) |
| NCBI nr/RefSeq protein | *Amaranthus tricolor* | AGL27 (96.26%) |
| | *Beta vulgaris* subsp. *maritima* | FLC-like 1 (59.82%) |
| | *Spinacia oleracea* | FLC-like (59.91%) |
| Swissprot | *Vitis vinifera* | AP1 (38.76%) |
| | *Oryza sativa* | MADS14 (39.89%) |
| | *Arabidopsis thaliana* | AGL70/FLC1-like (39.66%) |
| UniProtKb | *Spinacia oleracea* | FLC-like (60.4%) |
| | *Coffea arabica* | AGL27 (48.7%) |

(b)

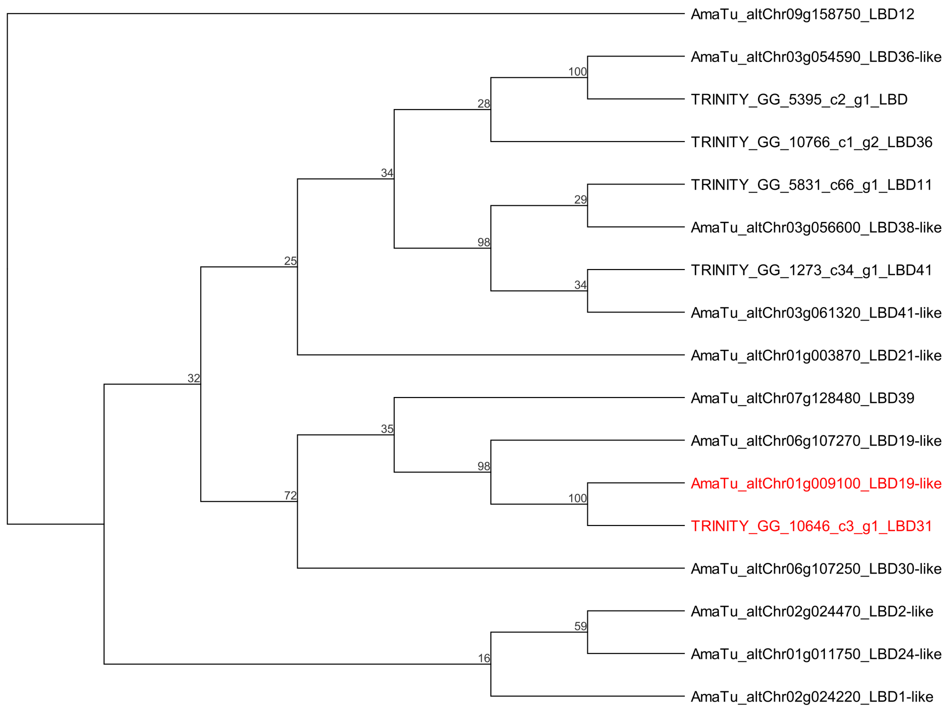


**Fig. S16** Alignment of MADS-box transcription factor 18 (MADS18) and MADS-box protein FLOWERING LOCUS C-like (FLC-like) amino acid sequence, and phylogenetic tree of LOB domain-containing protein (LBD). (a) Sequence alignment between MADS18 in Bobadilla *et al.*,

(2023) and FLC-like in this work. The table below indicates FLC-like annotations from different databases. (b) Phylogenetic tree of LBD. The tree leaves represent the various copies of LBD including copies from Bobadilla *et al.*, (2023). Leaves in red represent copies downregulated in males in that study and this work. Numbers above branches indicate bootstrap support values. Sequence alignment and tree were constructed using CLC sequence viewer 8.0.

**Methods S1** Library preparation, sequencing, assembly, and annotation.

**Waterhemp genome construction.** The waterhemp genome was constructed using a combination of Bionano Optical genome mapping and HiFi PacBio long sequencing as well as Hi-C Seq Illumina short read data. Bionano mapping was performed as previously described(Hufford *et al.*, 2021), with some modifications. Approximately 500 mg of young leaf tissue were fixed in 2% formaldehyde, chopped, homogenized, and filtered through 40 and 100 µm cell strainers, according to manufacturer's protocol using the Prep™ Plant Tissue DNA Isolation Kit (Bionano Genomics, San Diego CA). The resulting nuclei were resuspended and cleaned from debris and other solids using a series of low-speed centrifugations (100 x g) followed by centrifugation at 2,500 x g to concentrate and wash nuclei. Nuclei were resuspended, centrifugated again at 2,500 x g, and embedded in low melting point agarose. The resulting ultra-high molecular weight (uHMW) DNA was recovered from the agarose plug by melting it at 65 C, followed by incubation in agarase at 43 C and drop dialysis against TE buffer. The Bionano Direct Label and Stain (DLS) protocol was performed using an 800-ng uHMW with a DLS Kit, as per manufacturer's protocol (PN 80005), with some modifications. DNA was incubated in the presence of DLE-1 Enzyme, DL-Green and DLE-1 Buffer for 3 hours 40 minutes at 37 C and later incubated with Proteinase K for 1 hour at 50 C. After dye cleaning the reaction was stained, diluted in flow buffer, and loaded into a single Bionano chip in a Saphyr system as per manufacturers recommendations. Genome map assembly and hybrid scaffold construction were performed using Bionano Access version 1.7 and Tools 3.7 in a local Bionano Computer server. A subset of 1,537,123 molecules with a minimum of 150 Kb, N50 length of 276 kbp and combined length of 1,750 Gbp were assembled using the non-haplotype, no-CMPR-cut parameters without extend-split. The preliminary, non-phased assembly of the filtered molecules resulted in 54 genome optical maps with an N50 of 32.94 Mbp and total length of 1,267.17 Mbp.

For PacBio HiFi sequencing, DNA was isolated from approximately 1 g frozen leaf tissue using the Nucleobond HMW DNA kit (Macherey-Nagel) following the manufacturer's protocol. Purified DNA was sheared to a narrow size distribution of 10 to 20 kb with a Megaruptor 3 system (Diagenode) set at speed 34. The sheared DNA was used to construct a SMRTBell HiFi library using the Express Template Prep Kit 2.0 (PN: 101-853-100) as per manufacturers protocol. DNA fragments were size selected using the 15-20 kb High Pass 75E protocol in a

PippinHT System (Sage Science; PN: HPE-7510). The resulting SMRTbell library was bound to the sequencing polymerase enzyme and ran in two SMRTCells in a PacBio Sequel IIe sequencer, yielding a total of 4,585,400 HiFi reads with an average of 15,579 bp and totaling 71,435,938,804 bp.

One Hi-C Seq chromatin conformation capture library was constructed using the Phase Genomics Proximo system, as per manufacturer's protocol (Cat. KT3040; Phase Genomics, Seattle, WA). The library was constructed from chromatin crosslinked and isolated from 1 gram of frozen leaf tissue as per Manufacturer's protocol. The amplified library was sequenced in an Illumina NovaSeq 6000 system. The sequencing yielded 146,208,874 PE150 paired clusters for a total sequence length of 43,862,662,200 bp.

**Contig assembly, hybrid scaffolding and pseudomolecule construction.** The PacBio HiFi reads were assembled using Hifiasm v0.16.1 (Cheng *et al.*, 2021). The primary assembly was filtered for minimum length and coverage thresholds of 70 kbp and 20x, respectively. The resulting filtered assembly had 411 contigs with a total length of 1,267.172 Mbp and contig N50 of 32.943 Mbp. This combined contig set was aligned to a filtered BioNano map set using the Bionano Genomics Solve software (v3.7_03302022) with conflict resolution and sequence contig overlap trim enabled. The resulting 38 hybrid HiFi/DLS scaffolds with a total length of 1,198.106 Mbp were manually confirmed and curated. The 38 hybrid scaffolds consisted of 18 matching pairs, corresponding to two haplotypes. The matching pairs were identified by splitting scaffolds into 100 bp tags and aligned to all contigs with minimap2 v2-2.24-r1122 (Li, 2018) to find best matching pairs and orientation. Sequence polishing was performed on the hybrid scaffolds using alignment to PacBio HiFi corrected reads using minimap2 v2-2.24-r1122 (Li, 2018). Consensus errors were identified in the read alignments using mpileup and corrected. Non-scaffolded contigs were assigned to Chr00. Hi-C data was used for validation of the hybrid scaffolds as pseudomolecules using Juicer v0.7.0 (Durand *et al.*, 2016b). Visualization of Hi-C map contacts by Juicebox (Durand *et al.*, 2016a) confirmed haplotype resolution of the 38 pseudomolecules and corresponded to 16 pseudomolecules per haplotype.

**Genome annotation.** The two chromosome-level haplotype assemblies had the chromosomes sorted by size and were annotated individually. *De novo* transposable element families in the two assemblies were identified using RepeatModeler v.2.0.2 (Flynn *et al.*, 2020), and the assemblies were softmasked using RepeatMasker v. 4.1.2 (http://www.repeatmasker.org/RepeatMasker/) to generate an annotation of repeat elements and BEDtools v.2.30.0 (Quinlan & Hall, 2010) to softmask the genome using that annotation. The Iso-Seq reads were mapped to the softmasked genome using pbmm2 v.1.10.0. Redundant transcripts were collapsed to unique isoforms using IsoSeq3 v.3.8.2 (https://github.com/PacificBiosciences/IsoSeq). The final gene model predictions were made using MakerP v.1.0, utilizing the collapsed gene predictions from IsoSeq3 and all annotated protein sequences from *A. hypochondriacus* available on NCBI. Predicted small proteins (<12 amino acids) were filtered out and the remaining gene models were renumbered using a standard nomenclature. Functional annotation of the gene models was prescribed using a series of tools that predict protein location, function, homology. using MultiLoc2 v.1.0 (Blum *et al.*, 2009) was used to predict the subcellular localization of all proteins and whether or not they had a secretory sequence and/or transmembrane domains. InterProScan5 v.5.47-82.0 (Jones *et al.*, 2014) and Uniprot (Consortium, 2023) were used to predict protein domains within all the proteins and prescribe Interpro IDs, GOterms, Panther IDs, and Kegg Terms. MMSeqs2 v.4.1 (Steinegger & Söding, 2017) was used to search both the Uniref50 database from UniProt and a custom NCBI protein database for the nearest annotated homologue (i.e., best hit) to prescribe function by whole-protein homology. The custom NCBI protein database contained a manually curated list of proteins that are known herbicide targets. The best hit to the Uniref50 database was converted into the KEGG Orthology (KO) ID (Kanehisa *et al.*, 2017) and prescribed. The genome annotation (i.e., the gff) was modified to include relevant functional information in the notes column for each gene including the Interpro ID, GO IDs, and the closest known annotated protein.

Additional annotations were carried out to identify the transcription factors in the genome assemblies using PlantTFDB v5.0 (http://planttfdb.gao-lab.org/prediction.php)(Tian *et al.*, 2020). Disease resistance genes in the genome were also annotated following the Disease Resistance Analysis and Gene Orthology (DRAGO 3) pipeline in PRGdb v4 (https://github.com/sequentiabiotech/DRAGO-API) (Calle García *et al.*, 2022). Transfer RNA (tRNA) genes were predicted using tRNAscan-SE v2.0.12 (Chan *et al.*, 2021), an integrated part of the Maker pipeline. Other non-coding RNAs including microRNAs (miRNA), small nuclear RNAs (snRNA) and small nucleolar RNAs (snoRNA) were annotated using a similarity search in Infernal v1.1.5 (Nawrocki & Eddy, 2013) against the Rfam database 14.10 (Kalvari *et al.*,

2021). Ribosomal RNAs (rRNA), 18S and 28S, were annotated using RNAmmer v1.2 (Lagesen *et al.*, 2007) with parameters: -S euk -m lsu,ssu.

**Methods S2** Genome characteristics and repeat analysis.

The assembly quality and completeness for both haplotypes were accessed using the Benchmarking Universal Single-Copy Orthologs (BUSCOs) v4.0.2 (Simão *et al.*, 2015). Genome characteristics were computed using "agat_sp_statistics.pl" from AGAT Toolkit v1.0.0 (Dainat, 2022) and "bbstats.sh" from BBTools (Bushnell, 2014). Ortholog inference was carried out using protein-coding genes from both haplotypes and three chromosome-level assembly of amaranths (*A. hypochondriacus*, *A. cruentus* and *A. tricolor*) in OrthoFinder v2.5.5 (Emms & Kelly, 2019).

Species-specific repeats in the two haplotype assemblies were first identified using *de novo* approach with RepeatModeler v2.0.4 (Flynn *et al.*, 2020). The RepBase database (RepeatMaskerEdition-20181026) (Bao *et al.*, 2015) was then combined with Dfam3.2 database in RepeatMasker, and queried using the famdb.py utility in RepeatMasker to obtain 'viridiplantae' repeats library. A separate LTR structural analysis was also carried out. First, the genome assemblies were analyzed with LTR_harvest from genometools v1.6.0 using the parameters: -minlenltr 100 -maxlenltr 7000 -mintsd 4 -maxtsd 6 -motif TGCA -motifmis 1 -similar 85 -vic 10 -seed 20 -seqids yes (Ellinghaus *et al.*, 2008), and then analyzed with LTR_FINDER_parallel using default parameters (Ou & Jiang, 2019). The output obtained from both LTR_harvest and LTR_FINDER_parallel were combined and parsed to LTR_retriever v2.9.5 (Ou & Jiang, 2018) to obtain a non-redundant LTR library for each haplotype using the default parameters.

The substitution rate of 2.81e-9 synonymous substitutions per site per year for the Chenopodioideae plastid *rbcL* gene was also specified with -u parameter within LTR_retriever to estimate the insertion age of LTR elements. The three libraries (species-specific consensus library of repeats, viridiplantae repeats library and non-redundant LTR libraries) were combined for each of the haplomes. The combined libraries were clustered using CD-HIT-EST v4.8.1 (Li & Godzik, 2006) with parameters: -c 0.8 -G 0.8 -s 0.9 -aL 0.9 -aS 0.9 -M 0 -T 48 to reduce redundancy. The non-redundant libraries were then used to annotate repeats in the haplomes using RepeatMasker v4.1.5 with parameters: -s -gff -lib -e rmblast (http://www.repeatmasker.org/RepeatMasker/). Annotated repeats were summarized with RepeatMasker's "buildSummary.pl" script. The distribution of LTRs was estimated using the perl script "LTR_sum.pl" within LTR_retriever.

*De novo* quality of intergenic and repetitive sequence space of both haplotypes was also accessed using LTR assembly index (LAI) from the LTR retriever pipeline (Ou & Jiang, 2018; Ou *et al.*, 2018). Putative centromeric repeats in both haplotypes were identified using two complimentary approaches. The first approach was searching the assemblies for centromeric repeats using CentroMiner (Lin *et al.*, 2023), which uses tandem repeat finder (TRF)(Benson, 1999), BLAST clustering and TE annotation from EDTA (Ou *et al.*, 2019) to identify centromere candidates in the genome. The second approach was running StainedGlass (Vollger *et al.*, 2022) with default parameters (window = 5000 and mm_f = 10000) to determine sequence identity within tandem repeat arrays. Telomeric repeats were identified by searching the telomere repeat motif $(TTTAGGG)_n$ (where n = 4) against both haplotypes using BLASTN (Camacho *et al.*, 2009) with parameters: -task blastn-short -db haplotypes -query primers -outfmt 6 -out output.
